# Diffusive Flux Analysis of Tumor Vascular Permeability for 3-Helix-Micelles in Comparison to Other Nanoparticles

**DOI:** 10.1101/708263

**Authors:** Marc Lim, Vishnu Dharmaraj, Boying Gong, Ting Xu

**Author notes:** Corresponding author at: 381 Hearst Memorial Mining Building, UC Berkeley, Berkeley, CA 94720. **Declarations of interest**: none.

## Abstract

Understanding the complex interplay of factors affecting nanoparticle accumulation in solid tumors is a challenge that must be surmounted to develop effective cancer nanomedicine. The tumor microenvironment is unique in comparison to healthy tissue, possessing elevated interstitial pressure that limits convective transport, hence leaving diffusive processes to predominate especially in the tumor’s necrotic core. Certain tumor types such as glioblastoma multiforme and pancreatic tumor are known to be poorly permeable, making them less accessible for nanoparticle drug delivery. Evidence indicates that small and long-circulating nanoparticles are ideal for taking advantage of the enhanced permeability and retention effect, even in such intractable tumor models. Three-helix-micelle (3HM) self-assembled nanoparticles possess these characteristics, and previous studies have shown that 3HM can achieve more favorable tumor accumulation and penetration than liposomes in several tumor models. The reason for its superior performance had yet to be determined, and thus we sought to examine its passive transport into tumors. In this paper we present a simple mathematical model based on diffusive flux to describe particle accumulation in tumors with respect to particle plasma pharmacokinetics. Fitting the diffusive flux equation to in vivo particle tumor concentration yields the particle effective permeability value, which for 3HM is 2.31 ± 0.18 x 10^−8^ cm/s in U87MG glioblastoma and 4.25 ± 0.91 x 10^−8^ cm/s in HT29 colon cancer murine models. Applying this diffusive flux model to other nanoparticles reported in literature enables the effect of plasma half-life to be decoupled from particle permeability in influencing tumor accumulation, reinforced trends reported in the field regarding the impact of particle size, and provided a semi-quantitative means of comparing various tumor models. This work is also the first demonstration, to the best our knowledge, of extracting particle permeability values from bulk biodistribution data obtained via positron emission tomography, as opposed to laborious tumor optical window intravital microscopy experiments. As such the analysis provided here presents a simple and accessible tool to further enhance nanomedicine development.

Graphical abstract

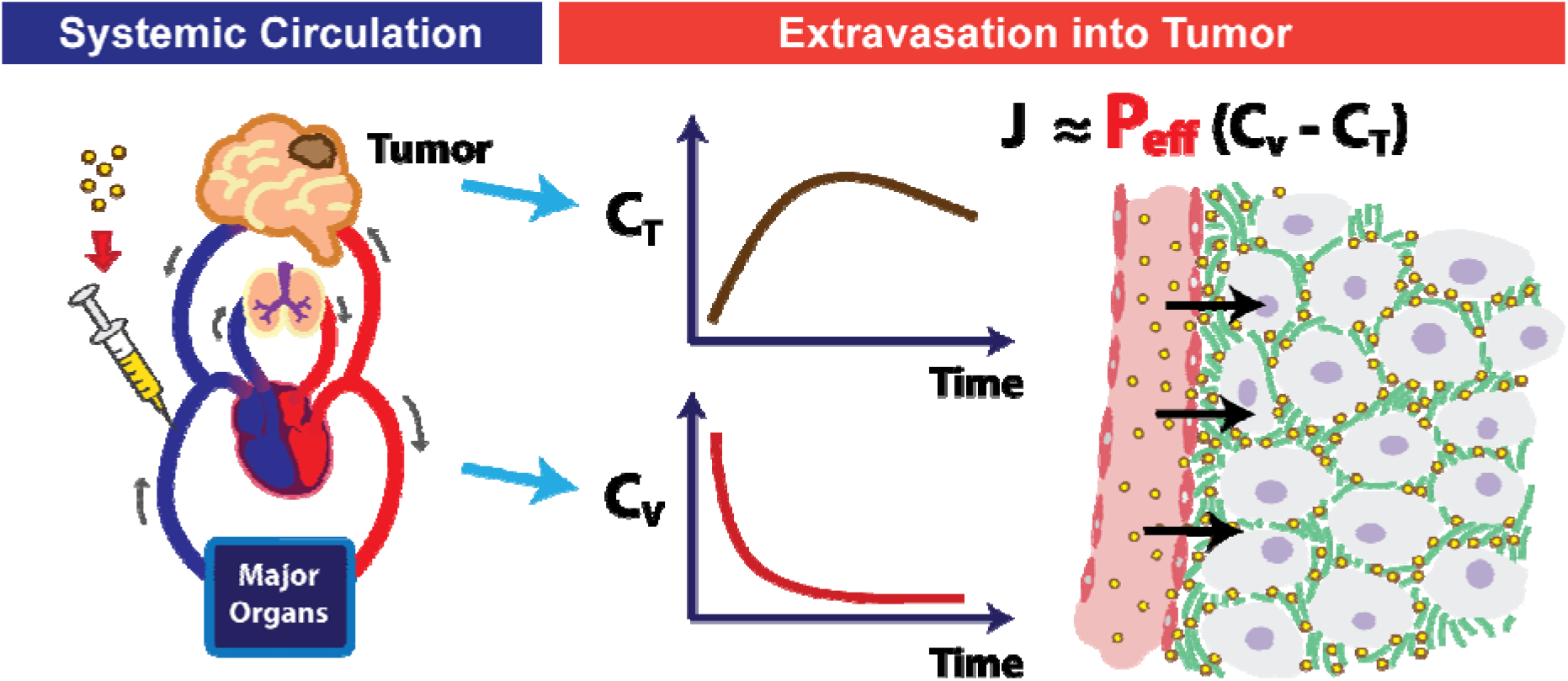

## Introduction

Cancer nanomedicine was built upon the premise that nanocarriers can improve their cargo’s pharmacokinetics, biodistribution, toxicity profile, and efficacy; however, lingering troubles have hindered these nanoparticles from fulfilling their promises [1,2,3]. In over forty years, various nanocarriers have been designed and tested to correlate cargo accumulation and antitumor activity with particle morphology, surface chemistry, material composition, mechanical properties, and the presence of active targeting ligands [4]. Yet, while thousands of papers have been published and hundreds of clinical trials have been attempted, only about a dozen cancer nanocarrier-based drugs have been approved worldwide [2,3,5]. The median delivery efficiency of nanocarriers remained stagnant at ~0.7% injected dose over the past decade, regardless of targeting strategy or material property [1]. While many studies are devoted to developing novel material synthesis, characterization, *in vitro* mechanistic exploration, and *in vivo* efficacy & biodistribution, few are devoted to studying the physiological transport processes governing nanoparticle accumulation and penetration through tumors. Such studies should be conducted with respect to the tumor microenvironment and a systemic-level perspective of the particle’s properties. A complex interplay of factors contributes to successful particle design, and without a definitive metric it is difficult to tease out which factor(s) predominates particle tumor transport. Having a model that could accomplish this will provide valuable knowledge for furthering the design and development of nanocarriers.

Numerous papers have delved into examining the tumor microenvironment, comprising of leaky vasculature and impaired lymphatic system that altogether allows extravasated nanoparticles to be entrapped in the tumor, leading to the enhanced permeability and retention (EPR) effect [6,7] [8]. Yet, despite having high vessel density and leaky vessel walls, particle transport towards the core of solid tumors remains poor and blood flow is heterogeneous [9]. This has been attributed to the high interstitial pressure at the necrotic core of solid tumors [10–13], which induces deformation and collapse of blood vessels at the tumor core, hence stunting fluid convection and leaving mostly diffusive processes to drive molecular transport [10]. Additionally, the size of tumor vessel fenestrations vary depending on tumor type, and while some such as MCaIV breast cancer can have very large fenestrations reaching up to 2 um, for U87MG glioblastoma brain tumors the fenestrations are on the order of 100 nm [14]. Naturally, the fenestrations create a size-exclusion effect for particles seeking to pass through, whereby particles greater than the pore size cutoff would be prevented from leaving the tumor vessels. Recent papers have highlighted the importance of small (<100 nm) and long-circulating particles (plasma half-life: t_1/2_ >6 hr) for effective tumor delivery, especially in poorly permeable tumors such as pancreatic cancer [15–17]. However, misconceptions still persist in the field regarding 100 nm being the optimal particle size for tumor accumulation [18,19], which arose from studies of liposomes of various sizes [20,21] and the precedent set by approved therapeutic liposome Doxil^®^. Likely, the optimal geometry of liposomes is dictated by their subunit packing parameter [22], such that extrusion into smaller diameters generated less stable structures that were cleared away faster upon injection. As such, the most stable form of the nanoparticle can vary greatly depending on its material, and it is important to select the right particle for successful performance in the intended application.

Several examples of promising sub-100 nm have been reported in literature, including block copolymer micelles, dendrimers, and chimeric polypeptide nanoparticles [23] [24] [25]. In addition, three-helix-micelle (3HM) nanocarriers developed in our lab has been found to achieve excellent tumor transport, with greater tumor accumulation and more favorable biodistribution than liposomes, even in an intractable cancer model such as orthotopic glioblastoma multiforme (GBM) [26–28]. 3HM is a self-assembled nanoparticle made up of peptide-polymer amphiphiles, containing alpha-helical coiled-coil head groups with hydrophobic alkyl tail core, polyethylene glycol (PEG) side-chains providing entropic repulsion, and PEGylated stealth outer layer [28,29]. 3HM is small (15-20 nm diameter) and yet has long plasma circulation (t_1/2_ = 29 hours in mice [28]) as compared to other nanoparticles such as liposomes (~100 nm diameter, t_1/2_ = 18 hours in mice [30]), and conventional surfactant micelles (7 – 20 nm diameter, t_1/2_ = 2 hours in mice [31]).

As such, we sought to develop a mathematical that could be applied broadly to nanoparticle tumor transport, with 3HM as a model particle, and surveyed the field to identify key transport parameters. Considering the lack of convective transport in tumors, diffusive flux becomes integral for particle movement into tumors. A concentration gradient drives particle extravasation from tumor blood vessels into the tumor interstitial space, which is aided by a long plasma t_1/2_. Another key parameter is particle permeability across vessel endothelium, whether transcellularly through endocytic pathways or intracellularly through vessel fenestrations, which determines the ease of particle extravasation for subsequent accumulation (Figure 1). Particle permeability has been thoroughly studied by Jain, Chilkoti, and Dewhirst’s groups via intravital microscopy of optical window tumor models [11,32–36]. However, such experiments require a very specialized skillset with limited availability to properly carry out, making such permeability analysis uncommon and under-utilized in nanoparticle development.

**Figure 1:**
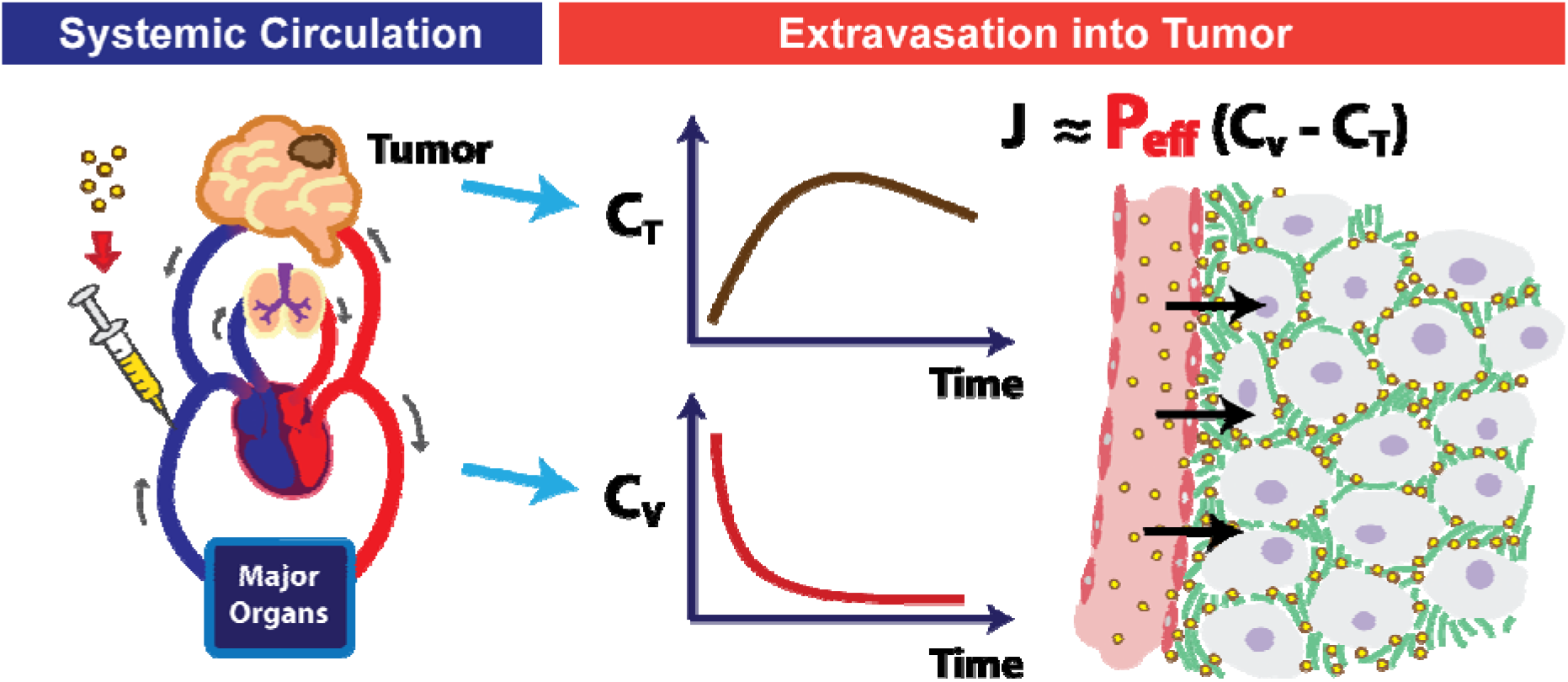
Schematic of intravenously administered particles migrating from systemic circulation to extravasation in a poorly-permeable brain tumor. Flux (J) of extravasated particles is dependent on flux area (A), particle permeability (P), and the concentration gradient in the tumor (C_T_) vs. blood vessel (C_v_).

Herein we present a simple mathematical model describing particle tumor concentration based on diffusive flux. The equation we derived matches what was reported by Jain, Stylianopoulos and colleagues [11] (refer to Supplementary Information for their paper), giving precedent to this analysis. What distinguishes our method is that this is the first instance, to the best of our knowledge, of correlating particle permeability to tumor accumulation using bulk biodistribution data (e.g. by positron emission tomography (PET) imaging) as opposed to via intravital microscopy. Fitting tumor concentration profile over time to this diffusive flux equation, along with particle plasma t_1/2_, allows particle permeability value to be derived for that specific tumor model. How well the accumulation data fits to the model can provide an indication as to whether diffusive, or convective, or other forces predominate for that particular system. We demonstrate this analysis with 3HM as a model nanoparticle, using previously reported PET data [26,37]. We then juxtapose 3HM’s results against various particle types and tumor models found in literature, using permeability values derived via a similar analysis. We believe that the trends presented here demonstrate the utility of using permeability values as a metric for informing particle design. While the diffusive flux model has some shortcomings compared to more elaborate models involving compartmental analysis [32] and convective fluxes arising from intratumoral pressure [11], its simplicity allows this model to be more widely accessible for many researchers seeking to develop effective cancer nanomedicine.

## Materials and Methods

### 1. Mathematical Model of Nanoparticle Flux

Nanoparticle tumor transport is governed by physical parameters, in addition to chemical and biological factors. Transvascular flux of particles crossing the tumor vessel endothelial wall barrier is described by the Staverman-Kedem-Katchalsky equation [38], which relies on both diffusive and convective fluxes:

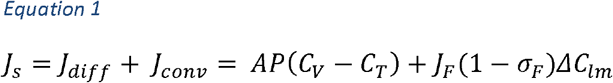

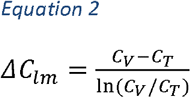

In Equation 1, *J*_*s*_ is the total solute (nanoparticle) flux (mg/s), *J*_*diff*_ is diffusive flux, *J*_*conv*_ is convective flux, and *J*_*F*_ is solvent fluid flux across the endothelial vessel wall (cm^3^/s). *A* is the flux area (cm^2^), *P* is the vascular permeability (cm/hr), *C*_*V*_ is particle concentration in the vessel (mg/cm^3^), *C*_*T*_ is particle concentration in the tumor interstitial spaces, *ΔC*_*lm*_ is the particle log mean concentration difference defined by Equation 2, and *σ*_*F*_ is the solvent-drag reflection coefficient (unitless). *J*_*F*_ is further described by the Starling Equation, which takes into account effects of hydrodynamic and oncotic pressures to determine migration of fluids from blood vessels to the interstitial spaces:

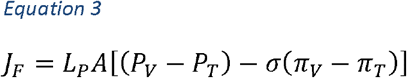

In Equation 3, *L*_*P*_ is the vessel hydraulic permeability (cm/s·mmHg), A is the vascular area for exchange (cm^2^) as above, *P*_*V*_ is the vessel hydrodynamic pressure (mmHg), *P*_*T*_ is the tumor interstitial fluid hydrodynamic pressure (mmHg), σ is the capillary reflection coefficient which describes the ease of solvent molecules crossing the vessel wall (unitless), *π*_*V*_ is the vessel colloid osmotic pressure (mmHg), and *π*_*T*_ is the tumor interstitial fluid colloid osmotic pressure (mmHg).

### 2. Diffusion-Governed Flux

The tumor microenvironment determines which of these factors are applicable for analysis. Direct measurement of *in vivo* tumor vascular and interstitial pressures have been attempted by several researchers, notably by Jain's group [13,39,40], but none of these measurements have been carried out in the same experiment as nanoparticle administration. It is well known, however, that the core of solid tumors have elevated pressure which hinder convective transport [10,11] such that diffusion chiefly governs particle movement in this case. As such, the convective term can be eliminated as follows:

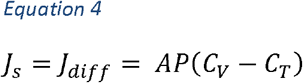

The migration of nanoparticles from blood vessels to tumor interstitial is schematically represented in Figure 2 as a 1-Dimensional diffusion flux problem. Here, flux *J*_*S*_ through a unit area, *A*, is examined, whereby the particle is expected to traverse a distance *L* through tumor tissue. *L* is dependent on the vascular density *S*_*V*_, which is the area of blood vessels per volume of tumor. Here *L* is assumed to be 100 um (0.01 cm) based on histology images of tumor vessels and *S*_*V*_ reported from literature [11].

**Figure 2:**
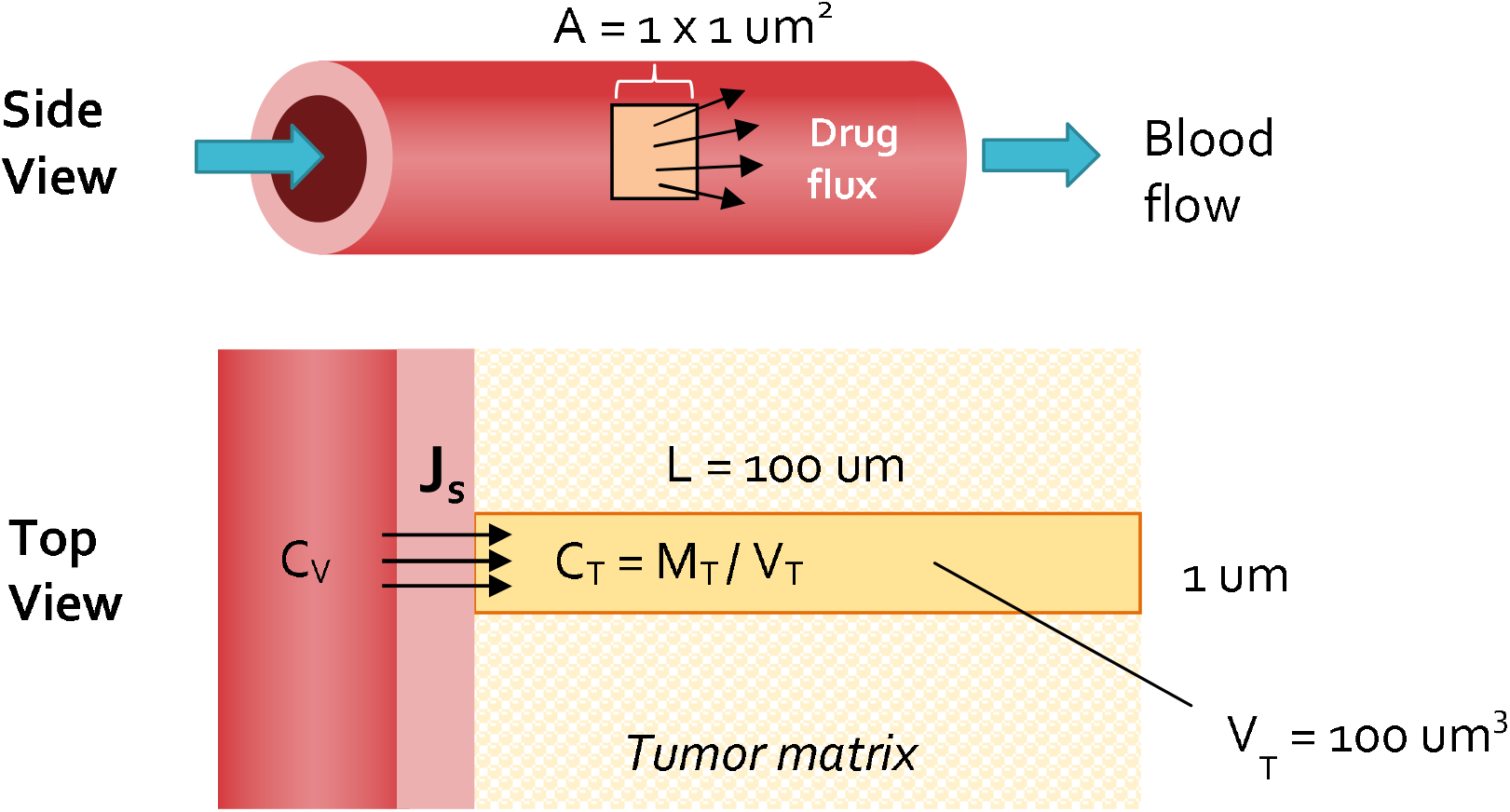
Schematic for modeling particle transvascular flux to tumor interstitia

Considering that particle concentration in the vessel, *C*_*V*_, decays exponentially over time due to elimination pathways (*C*_*V*_ = *C*_0_, *e*^−*Kt*^), the diffusive flux equation can be written to incorporate this term and solved as a first-order ordinary differential equation (Equation 5-Equation 7).

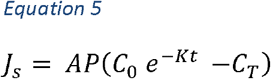

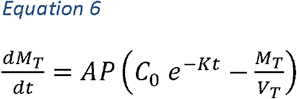

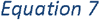

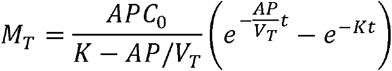

In Equation 5-Equation 7, *M*_*T*_ is the mass of the particle in the tumor, *C*_0_ is the initial particle concentration in the tumor vessel, *V*_*T*_ is the differential tumor volume being examined, and *K* is the particle plasma elimination time constant.

To obtain particle concentration in the tumor, particle mass *M*_*T*_ is divided by the tumor volume, *V*_*T*_:

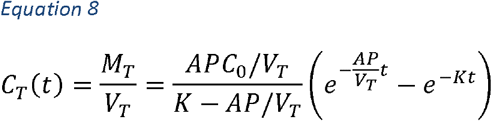

### 3. Fitting diffusive flux model to data from literature

Modelling data was extracted from articles found in the Cancer Nanomedicine Repository (http://inbs.med.utoronto.ca/cnr/) concerning the systemic delivery of particles to in vivo tumor models. For each particle and tumor model in an article, two sets of data were extracted as functions of time after injection: the particle concentration within the blood/plasma and the particle concentration within the tumor. The least squares exponential fit of plasma pharmacokinetics (PK) data (*C*_*V*_ = *C*_0_,*e*^−*Kt*^) was used to determine the particle plasma elimination time constant, *K*, and the initial particle concentration, *C*_0_, for each particle-tumor combination. After determining *K* for the system, empirical tumor accumulation data (*C*_*T*_) was fit to Equation 8 using the least squares method (Python’s SciPy “curve_fit” function). Since the permeability value is determined from empirical data using estimates, the result is termed as the effective vascular permeability, *P*_*eff*_. Values are reported with standard deviation of the fit. Equation 8 is therefore modified as follows:

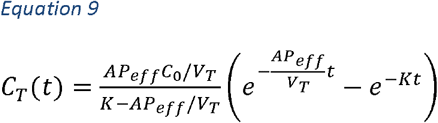

In this paper, the above analysis will be termed “**diffusive flux modeling**.”

Note: in practice, the concentration unit used for fitting can be presented in any form (e.g. %ID/g, mg/mL, mM), as long as it is kept consistent for both *C*_*T*_ and *C*_*0*_ since conversion factors would cancel out of both sides of Equation 9. Additionally, while most permeability values are reported in literature in cm/s, in this paper, permeability values are frequently presented in terms of um/hr as a more intuitive scale relating to distance in the tissue and cellular-level, and time-scale involved in drug pharmacokinetic studies. Unit conversions are presented where relevant for comparison purposes.

### 4. Statistical analysis

*P*_*eff*_ was estimated using least square fit of Equation 9. Under the assumption that Equation 9 gives unbiased estimations of the tumor accumulation and the normality of the errors, our estimated *P*_*eff*_ is normally distributed. Oftentimes, we are interested in comparing the *P*_*eff*_ fits between models. For example, when experiments are performed using different particles, we would be interested in finding the best particles with significantly larger permeability. Because of the nice normality property of our fitted *P*_*eff*_, two-sample t-test can be applied to compare between fits.

However, a few factors (see discussion in Results and Discussion 2.4) may introduce bias in our model. As a result, the estimated *P*_*eff*_ will be biased and so as the standard error estimation. Significance levels obtained using two-sample t-test would not be trustworthy. Standard regression diagnostic approaches such as residual plots can help to identify such biases. In the Supplementary Information, we discuss several fits and give examples where a two-sample t-test is or is not recommended.

## Results and Discussion

### 1. 3HM effective permeability in GBM and breast tumor models

Particle plasma PK (*C*_*V*_) and tumor accumulation data (*C*_*T*_) for 3HM and several other nanoparticles in NDL breast cancer and U87MG GBM orthotopic xenograft models [26,37] is presented in Figure 3 and Table 3. Based on estimates and available literature values for the other variables (*A* = 1 um^2^, *V*_*T*_ = 100 um^3^; see Figure 2), empirical *C*_*T*_ was fitted to the diffusive flux model described by Equation 9 using a Python algorithm, yielding particle-specific effective vascular permeability, *P*_*eff*_.

**Table 1:** Abbreviations

| Abbreviation | Term |
| --- | --- |
| 3HM | Three-helix-micelle |
| 3HM (C16) | Three-helix-micelle with palmitic acid tails |
| 3HM (C18) | Three-helix-micelle with stearic acid tails |
| 4T1 | Breast tumor lineage |
| Au NP | Gold nanoparticles |
| BCPM | Block-copolymer micelles |
| BSA | Bovine serum albumin |
| BxPC3 | Subcutaneous pancreatic cancer model |
| C26 | Subcutaneous colon cancer model |
| DOX | Doxorubicin |
| EAT | Ehrlich ascites tumor (breast cancer lineage) |
| EMT6 | Breast tumor xenograft lineage |
| EPR | enhanced permeability and retention |
| FaDu | Human squamous cell carcinoma lineage |
| GBM | Glioblastoma multiforme |
| HT29 | Colon tumor xenograft lineage |
| IgG | Immunoglobulin G |
| LS174T | Colon tumor xenograft lineage |
| MCalV | Breast cancer line |
| MET1 | Breast tumor lineage |
| NDL | Breast tumor xenograft lineage |
| NidoC | Nidocarborane |
| NS | Non-significant |
| PEG | Polyethylene glycol |
| PEI | Polyethyleneimine |
| PET | Positron emission tomography |
| PK | Pharmacokinetics |
| R3230AC | Mammary tumor xenograft lineage |
| Rho-BSA | Rhodamine-conjugated BSA |
| SCID | Severe combined immune deficiency |
| subQ | Subcutaneous |
| TRITC-BSA | Tetramethylrhodamine-conjugated BSA |
| U87MG | Glioblastoma xenograft lineage |
| VEGF | Vascular endothelial growth factor |
| Zr-CLL | Liposomes radiolabeled with Zr<sup>89</sup> using click-chemistry |
| Zr-SCL | Liposomes radiolabeled with Zr<sup>89</sup> using chelator |

**Table 2:** Mathematical variables

| Variable | Description |
| --- | --- |
| $A$ | Flux area |
| $C_0$ | Initial concentration in vessel |
| $C_T$ | Concentration in tumor |
| $C_V$ | Concentration in vessel |
| $\Delta C_{lm}$ | Particle log mean concentration |
| $J_F$ | Solvent fluid flux |
| $J_{conv}$ | Convective flux |
| $J_{diff}$ | Diffusive flux |
| $J_s$ | Total solute flux |
| $K$ | Particle plasma elimination time constant |
| $L$ | Particle migration distance |
| $L_P$ | Vessel hydraulic permeability |
| $M_T$ | Mass of particle in tumor |
| $P$ | Vascular permeability |
| $P_T$ | Tumor interstitial fluid hydrodynamic pressure |
| $P_V$ | Vessel hydrodynamic pressure |
| $P_{app}$ | Apparent permeability, analogous to $P_{eff}$ |
| $P_{eff}$ | Effective vascular permeability |
| $S_V$ | Tumor vessel density |
| $V_T$ | Tumor volume |
| $t_{1/2}$ | Half-life |
| $\pi_T$ | Tumor interstitial colloid osmotic pressure |
| $\pi_V$ | Vessel colloid osmotic pressure |
| $\sigma_F$ | Solvent-drag reflection coefficient |
| $\sigma$ | Capillary reflection coefficient |

**Table 3:**
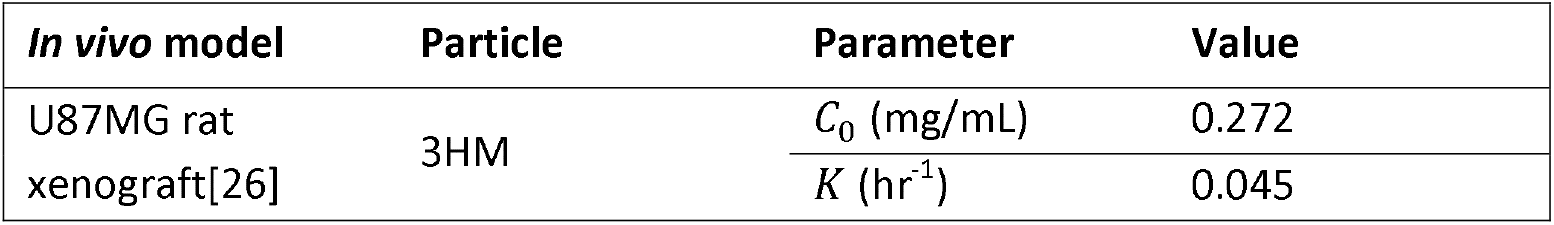
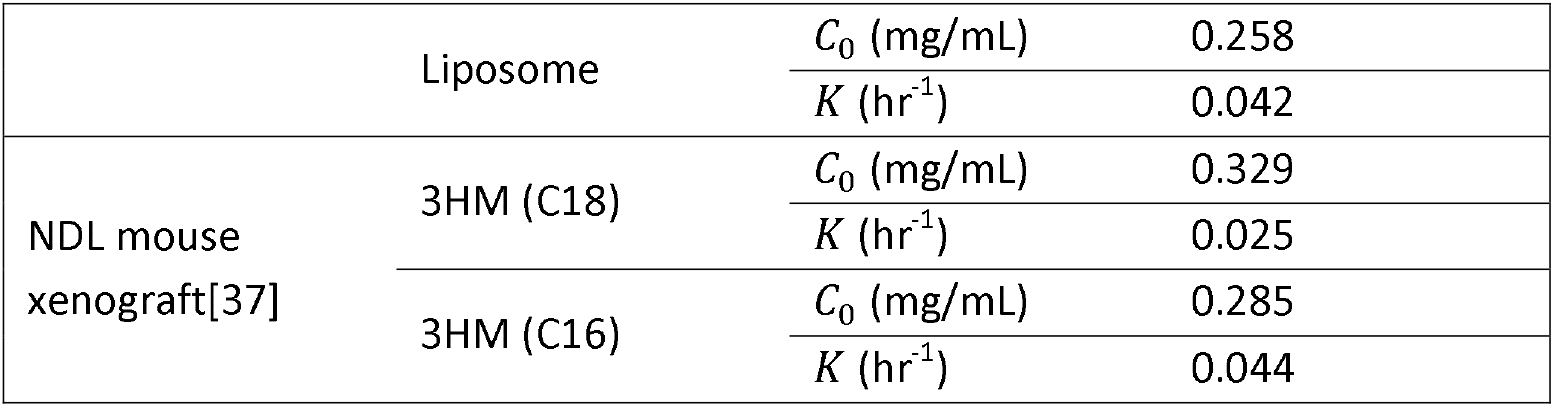
Parameter values for 3HM and Liposome plasma pharmacokinetics, *C*_*v*_ = *C*_*0*_ exp (−Kt)

**Figure 3:**
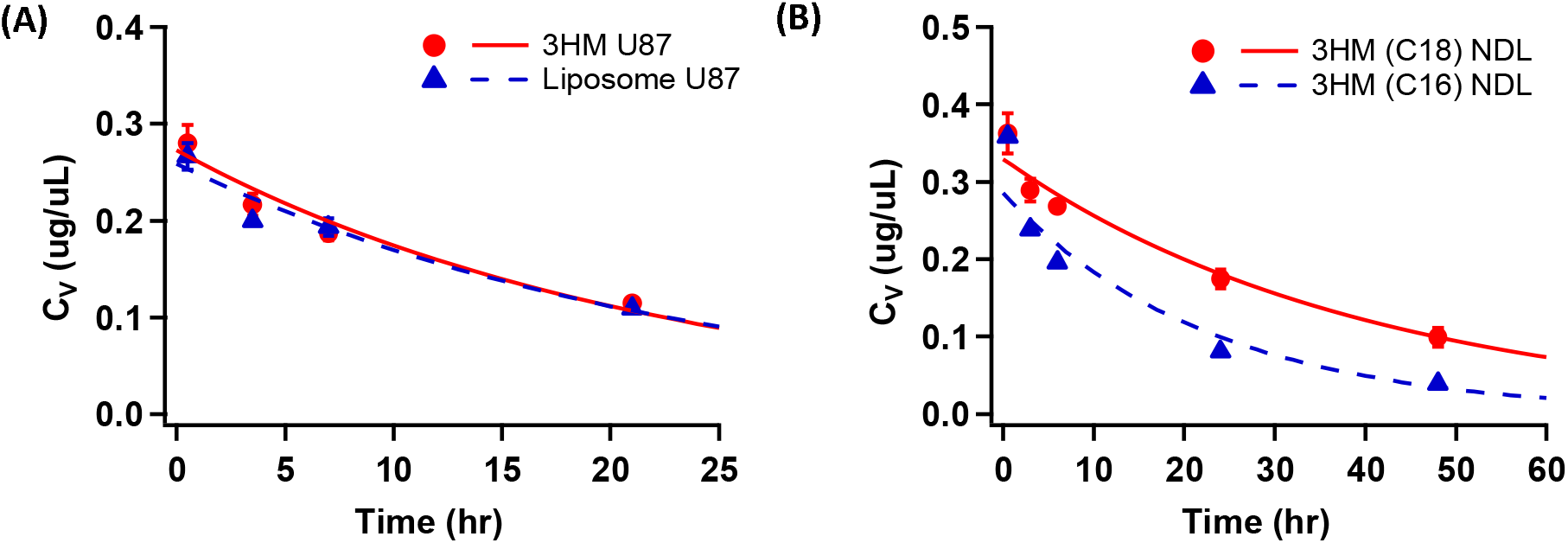
Plasma PK profiles of 3HM vs. other particles in (A) U87MG glioblastoma[26] and (B) NDL breast tumor[37] orthotopic in vivo models. Markers show empirical data and lines show trendline fit to a single exponential model (3HM U87MG: 0.272 exp(−0.0446t); Liposome U87MG: 0.258 exp(−0.0421t)); 3HM(C18) NDL: 0.329 exp(−0.025t); 3HM(C16) NDL: 0.285 exp(−0.044t))

In the U87MG GBM model, 3HM is compared to liposomes, while in the NDL breast tumor model, 3HM with two different amphiphile architectures are compared: ones with stearic acid alkyl tails (C18) customarily used for 3HM preparation, versus shorter palmitic acid tails (C16). The results presented in Figure 4 show that the model fits very well for U87MG brain tumor but poorly for NDL breast tumor data. This is not surprising considering diffusive transport governs macromolecular transport in the brain [41]. The poor fit in NDL data set indicates that other factors such as convective transport may be more dominant in this model, and a revised model would be necessary to fully explain the data.

**Figure 4:**
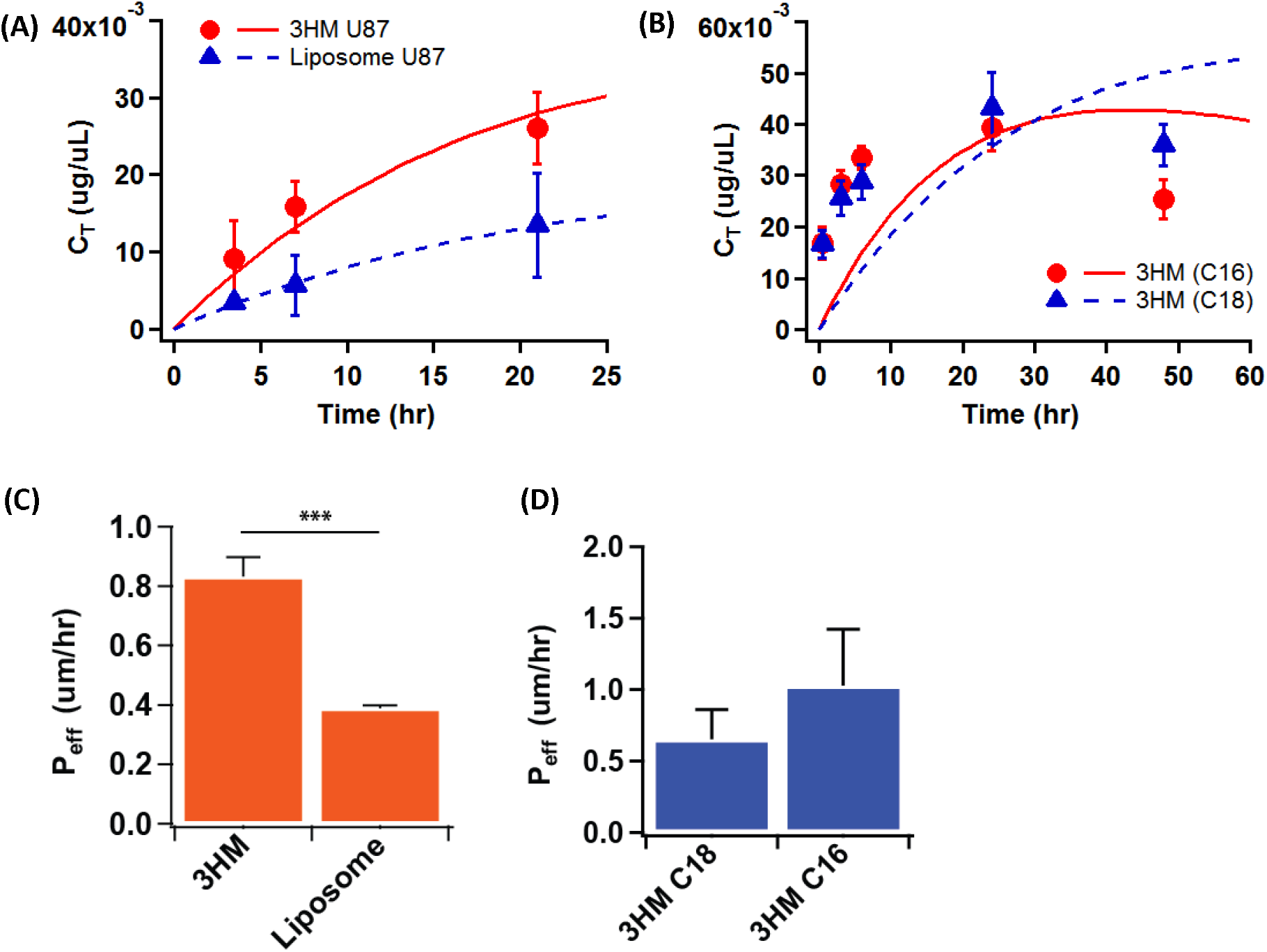
Nanoparticle tumor accumulation profiles in (A) orthotopic U87MG GBM and (B) orthotopic NDL breast tumor xenograft. Markers show empirical data and lines show trendline fit to diffusion flux model. P_eff_ values from model fit are compared in (C) for U87MG brain tumor, and (D) for NDL breast tumor. Error bars represent standard deviation. (***P < 0.0001 by Student’s t-test)

### 2. Comparison of diffusive flux modeling to other published methods of measuring particle permeability

To determine the utility and performance of our methods, *P*_*eff*_ of Dextran obtained by diffusive flux modeling was compared to values reported by Chilkoti and colleagues (termed “apparent permeability” or *P*_*app*_ in their paper) [32]. The results are shown in Figure 5 and Table 4.

**Table 4:** Dextran P_eff_ fit obtained using the diffusion flux model vs. those reported in literature [32]. Dextran hydrodynamic diameters (D_H_) are approximated using the Stokes-Einstein equation.

| Dextran | Approx. $D_H$<br>(nm) | $P_{eff}$ from current model (A) | | Median $P_{eff}$ in<br>literature[32] (B) | Ratio<br>of (A) /<br>(B) |
| --- | --- | --- | --- | --- | --- |
| | | (um/hr) | (cm/s) x $10^{-7}$ | (cm/s) x $10^{-7}$ | |
| 2 Mda | ~ 50 | 34.91 ± 0.76 | 9.70 ± 0.21 | 1.7 | 5.7 |
| 70 kDa | ~ 15 | 202.0 ± 21.3 | 56.1 ± 5.8 | 9.8 | 5.7 |
| 40 kDa | ~ 11 | 236.7 ± 18.2 | 65.8 ± 5.1 | 9.5 | 6.9 |
| 10 kDa | ~ 5 | 526.5 ± 136.3 | 146 ± 37.9 | 32 | 4.6 |
| 3.3 kDa | ~ 3 | 9102 ± 2296 | 2528 ± 638 | 154 | 16.4 |

**Figure 5:**
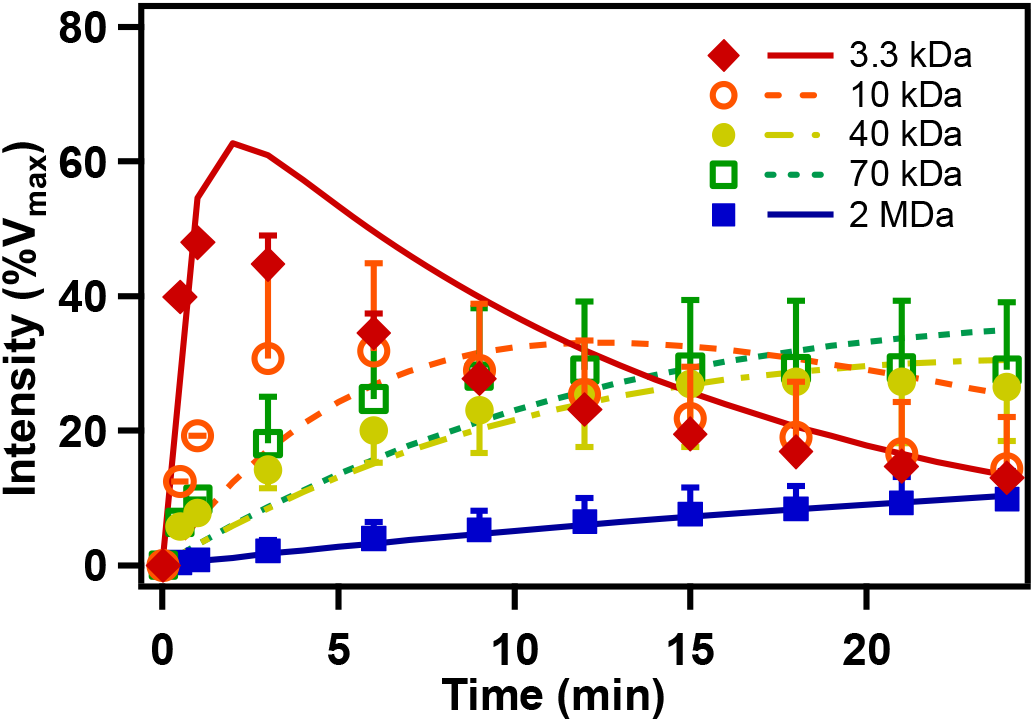
Accumulation of fluorescently-labeled 3.3 kDa – 2 MDa Dextran in human squamous cell carcinoma (FaDu) in vivo cranial window tumor model, presented as percent of maximum vascular intensity recorded. Markers refer to data points gleaned from literature [32], while trendlines show fits obtained using diffusion flux model.

Our model is notably simpler: (1) the plasma PK profile is approximated by a simple exponential function, as opposed to the bi-exponential curve typical for particles with alpha and beta-phase elimination half-lives; (2) it does not take into account hematocrit volume fraction in blood vessels, or particles volume fraction in tumor extravascular space; (3) the overall tumor accumulation is not fitted to a four-compartmental analysis model.

We considered Simplification (1) to be reasonable since the alpha-phase PK of particle distribution occurs very rapidly, and therefore contribute little to overall particle accumulation in tumors. Simplification (2) and (3) likely led to the roughly 6x difference between *P*_*eff*_ obtained by our model vs. those reported by the paper, for everything except the smallest Dextran particle (see Table 4).

This exercise has shown that a simple and fast diffusion flux modeling method could be used to compare *P*_*eff*_ of various particles within the same experimental tumor model. Except for the smallest particle tested (3.3 kDa, ~3 nm hydrodynamic diameter (D_H_)), the relative differences in dextran *P*_*eff*_ with respect to molecular weights are comparable to those reported by Chilkoti and colleagues, differing consistently by a factor of ~6.

Additionally, *P*_*eff*_ values for 3HM and liposomes obtained by our diffusive flux model can be compared to values obtained by Dr. Rakesh Jain, Dr. Mark Dewhirst and colleagues using fluorescently labeled liposomes and bovine serum albumin (BSA) particles, imaged by intravital microscopy (Table 5).

**Table 5:**
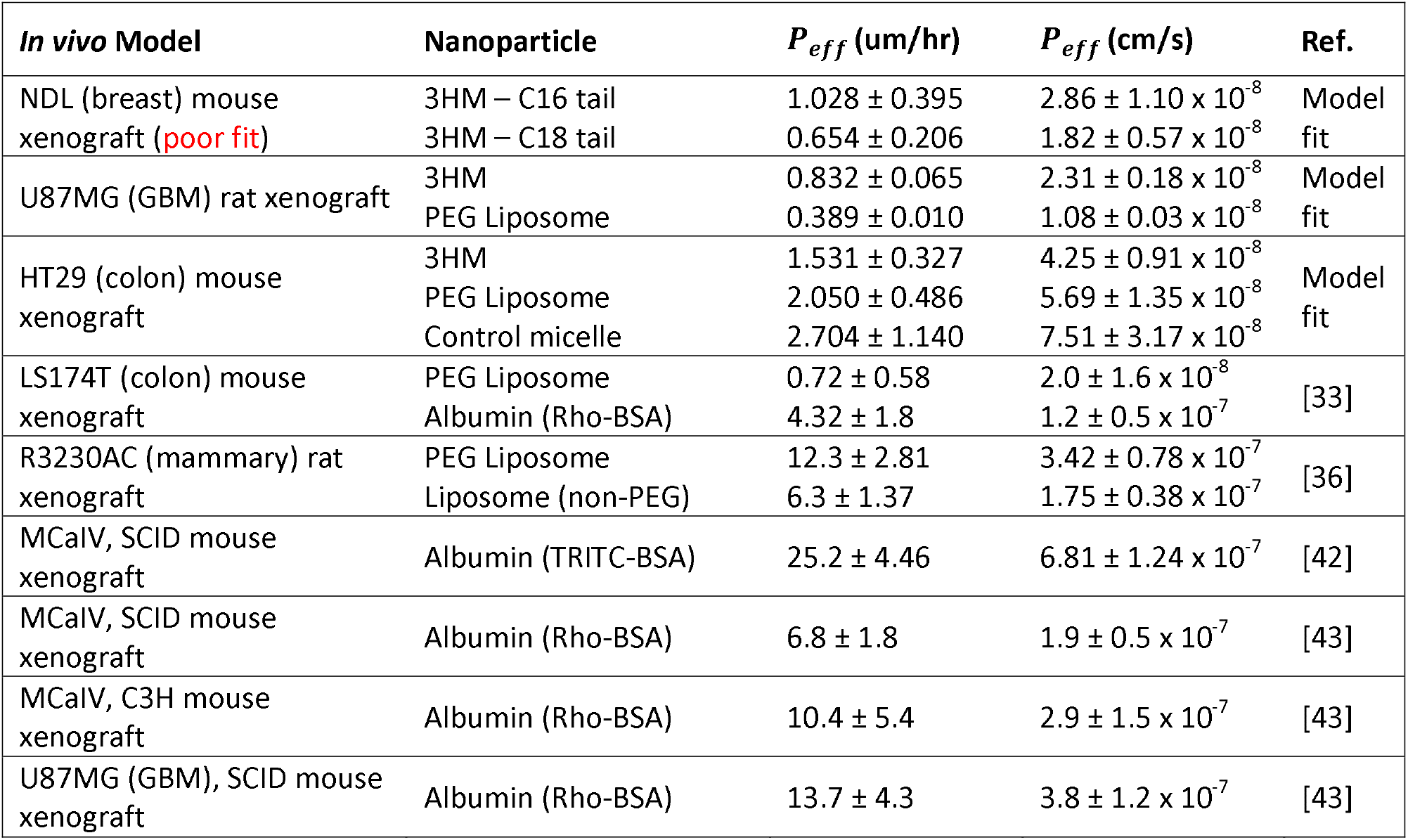
Comparison of P_eff_ values between mathematical model fits and literature values. Unless otherwise noted, all liposomes listed below refer to PEGylated “stealth” liposomes

| **In vivo Model** | **Nanoparticle** | **$P_{eff}$ (um/hr)** | **$P_{eff}$ (cm/s)** | **Ref.** |
| --- | --- | --- | --- | --- |
| NDL (breast) mouse xenograft (poor fit) | 3HM – C16 tail | 1.028 ± 0.395 | 2.86 ± 1.10 × 10<sup>-8</sup> | Model fit |
|  | 3HM – C18 tail | 0.654 ± 0.206 | 1.82 ± 0.57 × 10<sup>-8</sup> |  |
| U87MG (GBM) rat xenograft | 3HM | 0.832 ± 0.065 | 2.31 ± 0.18 × 10<sup>-8</sup> | Model fit |
|  | PEG Liposome | 0.389 ± 0.010 | 1.08 ± 0.03 × 10<sup>-8</sup> |  |
| HT29 (colon) mouse xenograft | 3HM | 1.531 ± 0.327 | 4.25 ± 0.91 × 10<sup>-8</sup> | Model fit |
|  | PEG Liposome | 2.050 ± 0.486 | 5.69 ± 1.35 × 10<sup>-8</sup> |  |
|  | Control micelle | 2.704 ± 1.140 | 7.51 ± 3.17 × 10<sup>-8</sup> |  |
| LS174T (colon) mouse xenograft | PEG Liposome | 0.72 ± 0.58 | 2.0 ± 1.6 × 10<sup>-8</sup> | [33] |
|  | Albumin (Rho-BSA) | 4.32 ± 1.8 | 1.2 ± 0.5 × 10<sup>-7</sup> |  |
| R3230AC (mammary) rat xenograft | PEG Liposome | 12.3 ± 2.81 | 3.42 ± 0.78 × 10<sup>-7</sup> | [36] |
|  | Liposome (non-PEG) | 6.3 ± 1.37 | 1.75 ± 0.38 × 10<sup>-7</sup> |  |
| MCalV, SCID mouse xenograft | Albumin (TRITC-BSA) | 25.2 ± 4.46 | 6.81 ± 1.24 × 10<sup>-7</sup> | [42] |
| MCalV, SCID mouse xenograft | Albumin (Rho-BSA) | 6.8 ± 1.8 | 1.9 ± 0.5 × 10<sup>-7</sup> | [43] |
| MCalV, C3H mouse xenograft | Albumin (Rho-BSA) | 10.4 ± 5.4 | 2.9 ± 1.5 × 10<sup>-7</sup> | [43] |
| U87MG (GBM), SCID mouse xenograft | Albumin (Rho-BSA) | 13.7 ± 4.3 | 3.8 ± 1.2 × 10<sup>-7</sup> | [43] |

It is apparent that while there are various *P*_*eff*_ values across different tumor models and particle types, the diffusive flux model fitted values reported here lie within an order of magnitude to literature values for liposomes. 3HM being a smaller particle (~15-20 nm [26,44]) has a higher *P*_*eff*_ than liposomes (~100 nm [26]), akin to results in colon cancer xenograft model for liposomes vs. BSA (~8 nm [45]).

Interestingly, literature values also demonstrate the wide variability of *P*_*eff*_ for albumin, even for the same tumor type, evidenced by TRITC-BSA being nearly 4x greater than Rho-BSA values in MCaIV xenografts [42,43]. It is unlikely that the conjugated fluorophores influenced this effect to such a degree, and thus is more likely indicative of tumor heterogeneity [46] as well as inherent variability of *in vivo* models. This variability has also been reported for albumin tested in rats with R3230Ac breast tumors (N=14), where *P*_*eff*_ ranged from 2-16 x 10^−7^ cm/sec [47].

### 3. Permeability values fitted for various nanoparticles

Accordingly, the diffusion flux modeling method described above was applied to a spread of nanoparticle biodistribution results reported in literature, to obtain their relative *P*_*eff*_ values. A total of 63 papers were perused for this effort, obtained mostly from the Cancer Nanomedicine Repository (http://inbs.med.utoronto.ca/cnr) maintained by Warren Chan and colleagues, which was borne out of an extensive review of nanoparticle delivery efficiency to tumors[1]. From that pool, 37 appeared suitable for analysis (i.e. reported data contains both plasma pharmacokinetics and tumor accumulation data, with at least three time-points examined), but only 19 were able to achieve reasonable fits (reasons for poor fits discussed below). Details of particle types, tumor models, fitting parameters and results are shown in Table S1.

The following *P*_*eff*_ trends emerged when comparing particles across various tumor types, which corroborates findings generally known in the field of nanoparticle delivery:

1. *P*_*eff*_ values for breast cancer > colon > pancreatic ≈ GBM
2. Larger particles have lower *P*_*eff*_ in poorly permeable tumors
3. For two particles with the same *P*_*eff*_, longer plasma circulation leads to more tumor accumulation

We had also examined the effect of active targeting on particle *P*_*eff*_, yielding some trends nm particles in various depending on the targeting ligand used (see Supplementary Information). However, more in-depth analysis would have to be performed before any conclusions can be made in this regard.

#### 3.1. Permeability of ~100 nm particles across various tumor models

To compare the *P*_*eff*_ of various tumors, we analyzed tumor accumulation data of ~100 nm particles found in literature according to the diffusion flux model. This particle size was chosen due to the precedent set by FDA-approved therapeutic liposomes (Doxil^®^, Marqibo^®^, Onivyde^®^, Ambisome^®^, and Amphotec^®^) which are similarly-sized [5]. The *P*_*eff*_ value comparisons are shown in Figure 6. The tumor xenograft cell lines are listed, along with whether they were subcutaneously (subQ) or orthotopically inoculated. The particles compared comprised of polyethyleneimine branched dendrimers (PEI branch) [48], liposomes [26,49–52], and block-copolymer micelles (BCPM) [16]. For the colon cancer data set, liposomes encapsulated either nidocarborane (NidoC) [51] or doxorubicin (DOX) [52] payloads.

**Figure 6:**
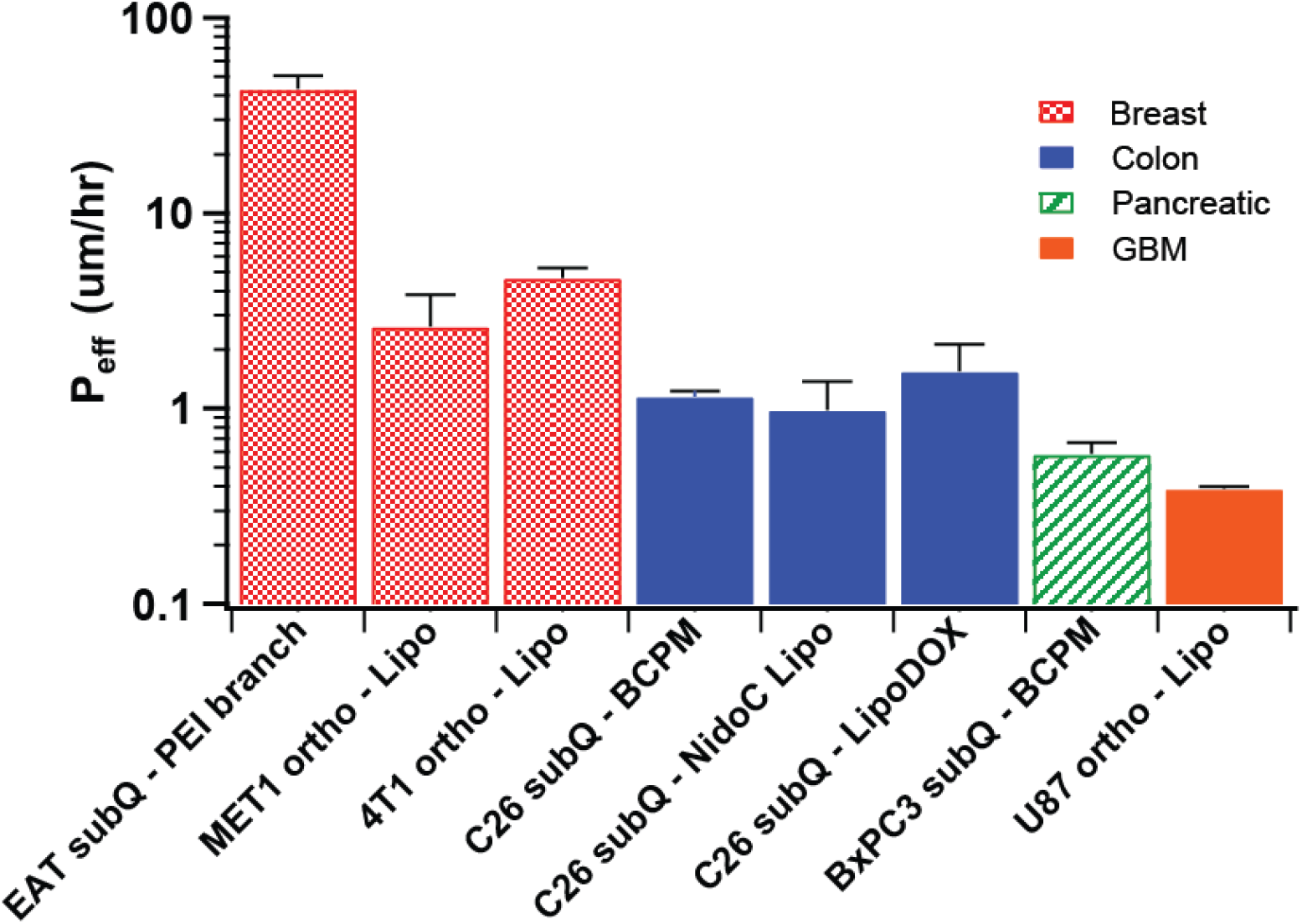
P_eff_ of ~100 nm particles in various murine tumor models. “Lipo” refers to liposomes and “ortho” for orthotopic inoculation; all other abbreviations are explained in the text body.

In Figure 6, a general trend emerged for *P*_*eff*_ among various tumors, with breast tumors being the most permeable, followed by colon cancer, pancreatic, and GBM. This corroborates other findings literature describing pancreatic and GBM tumors to be among the least permeable tumors [14,16,53] Note that liposome *P*_*eff*_ values in breast and colon tumors measured by other methods, shown in Table. 5, also fall within their respective ranges shown in Figure 6.

The *P*_*eff*_ of Ehrlich ascites tumor (EAT) breast cancer dosed with PEI dendrimer is an order of magnitude greater than for the other two breast cancer models, which could potentially be a result of the xenograft cell lineage used (i.e. EAT tumors being more permeable than MET1 or 4T1 tumors), or the tumor inoculation location (i.e. subcutaneous flank breast tumors are more permeable than orthotopic tumors in mammary fat pads). When the *P*_*eff*_ values of tumors with similar lineage and inoculation location were compared, namely for the C26 subcutaneous colon cancer models, the values were similar. This indicates that inter-experiment comparison is possible and that C26 subcutaneous tumors have reproducibly consistent permeabilities to ~100 nm particles even across different particle types and experimental research groups.

#### 3.2. Effect of particle size in various tumor models

Kataoka and colleagues have published an article detailing the differences in tumor accumulation and penetration of 30 – 90 nm block copolymer micelles [16], observed by intravital microscopy. The *P*_*eff*_ values for these experiments fitted by our diffusion flux model is shown in Figure 7A and B.

**Figure 7:**
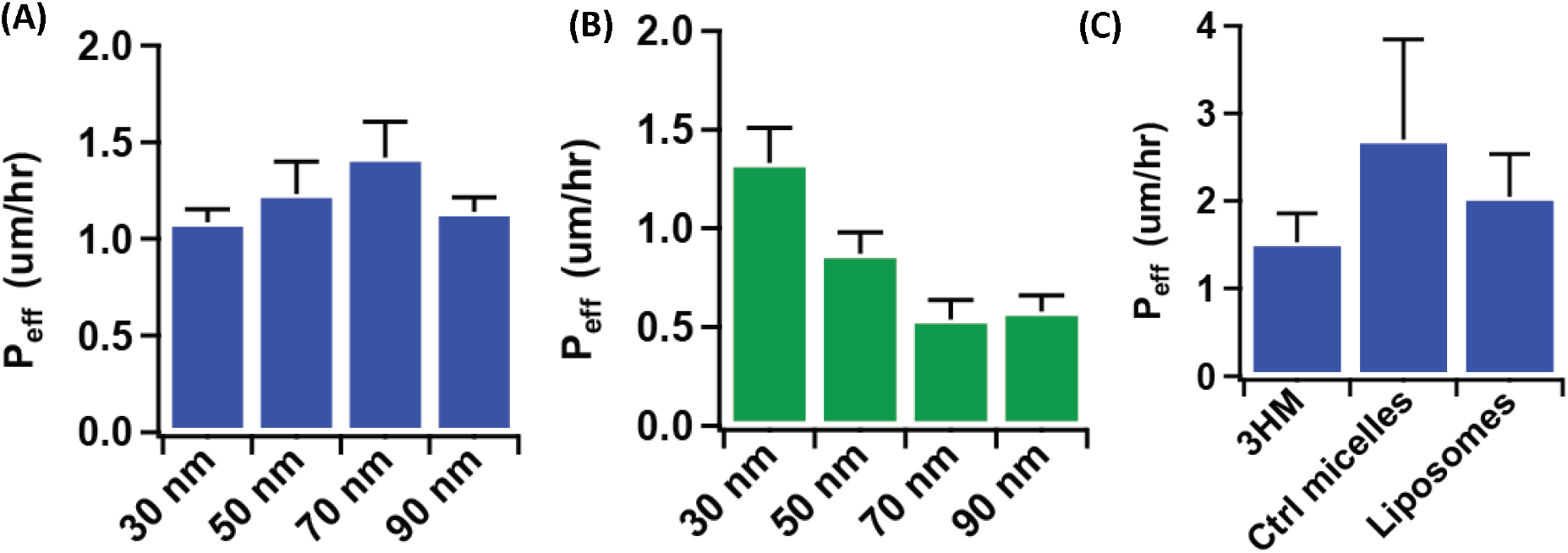
P_eff_ of 30-90 nm BCP micelles in (A) C26 subcutaneous colon cancer model and (B) BxPC3 subcutaneous pancreatic cancer model. (C) P_eff_ of 3HM (~20 nm), control micelles (~20 nm), and liposomes (~100 nm) in HT29 colon tumor.

It is apparent that in the colon cancer model, the effect of particle size was largely insignificant, while for pancreatic cancer model, there is a decrease in *P*_*eff*_ for larger particles that demonstrate a size-exclusion effect. Interestingly, *P*_*eff*_ for all particles in the colon cancer model are approximately similar to 30 nm particles in pancreatic cancer, hence only the smallest particle tested in pancreatic tumor was able to achieve the same level of permeability as in the colon tumor. The lack of a correlation between permeability to particle size (100 nm and below) in colon tumors was also observed in an experiment comparing 3HM (~20 nm), conventional micelles (~20 nm), and liposomes (~100 nm) in a HT29 colon cancer mouse model (Figure 7C). The conventional micelles accumulated least in the tumors compared to the other two particles, and based on their relative *P*_*eff*_ value, this is likely due to its shorter plasma t_1/2_ of conventional micelles (data not shown).

The effect of particle size on diffusive flux-modeled *P*_*eff*_ in various tumors are compiled in Figure 8, including 3HM and liposome data in GBM shown previously (Figure 4). Breast tumor *P*_*eff*_ values have been excluded considering that diffusive modeling may not be suitable for this tumor type. Figure 4 summarizes that particle permeability is generally independent of size for colon tumors, but dependence can be observed for pancreatic and GBM tumors.

**Figure 8:**
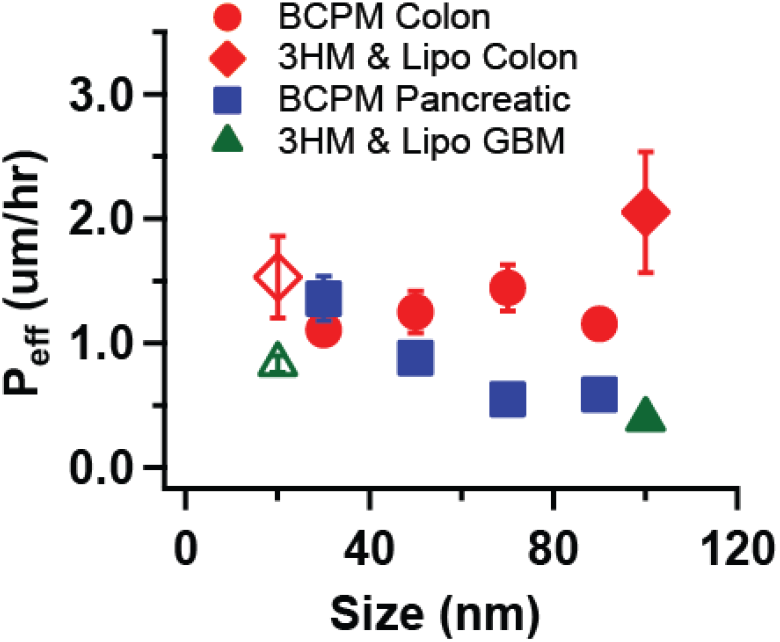
Effect of particle size on P_eff_ for various tumor models. BCPM = block copolymer micelles, Lipo = liposomes. Open symbols represent 3HM data points.

### 4. Decoupling effect of plasma PK from particle permeability in driving tumor accumulation

In some cases, it can be difficult to determine which factors impact tumor accumulation when modifying a particle property. Consider the following hypothetical scenarios (Figure 9), whereby two particles have different tumor accumulation profiles. In Figure 9A, since both particles have similar PK profiles, one can intuitively attribute differences in tumor accumulation to differences in *P*_*eff*_. In Figure 9B and C, it is less obvious, since in each of these cases the particles have different PK profiles. It is only upon fitting the data that it becomes clear that the particles in Figure 9B actually have similar *P*_*eff*_ values, while those in Figure 9C have different *P*_*eff*_ values.

**Figure 9:**
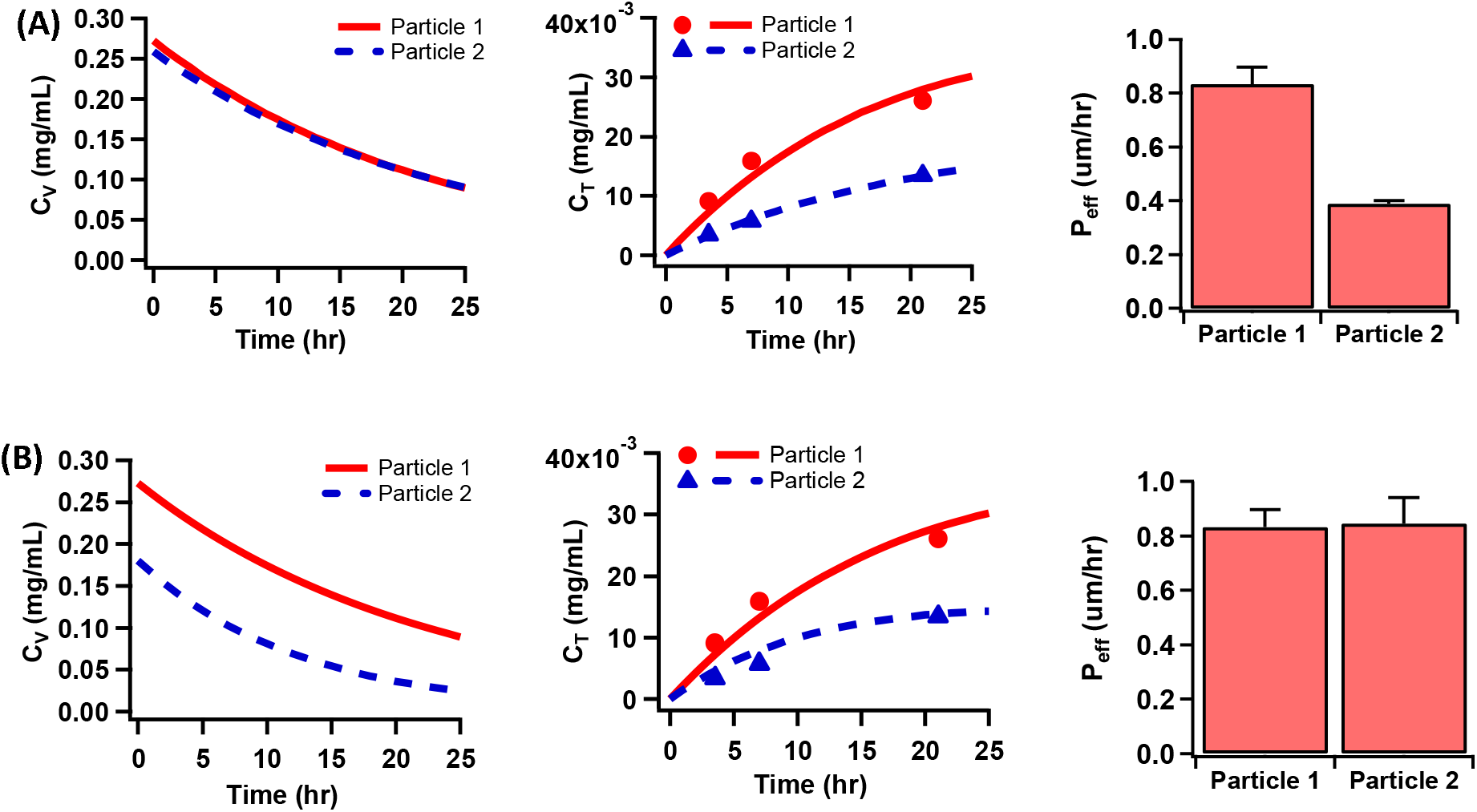
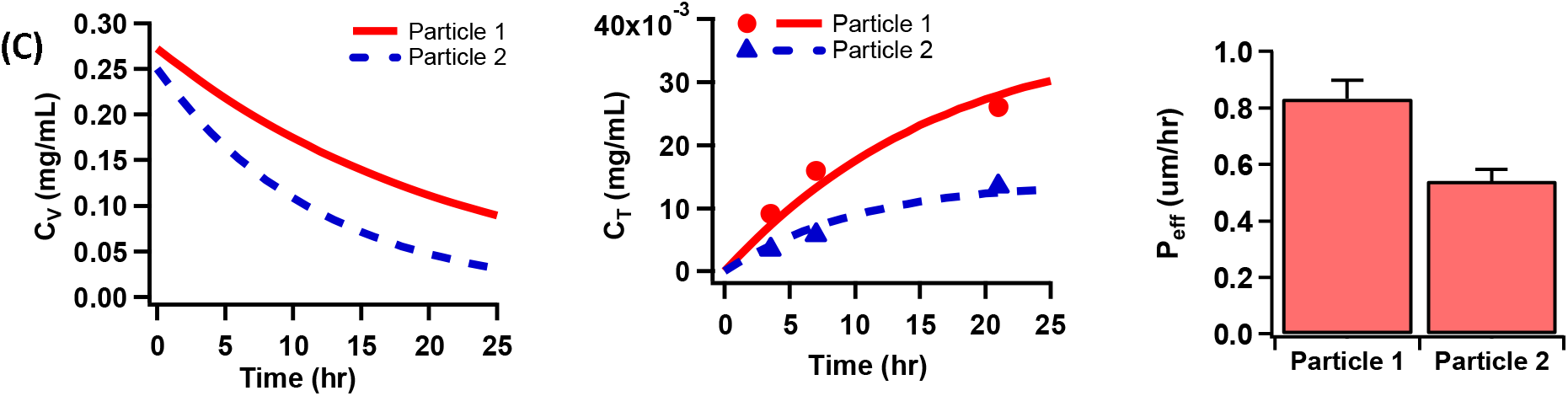
Hypothetical scenarios of two particles with different tumor accumulation: (A) Having similar PK profiles, but different P_eff_. (B) Having different PK profiles, but similar P_eff_. (C) Having different PK profiles and P_eff_ values.

Such insights can prove valuable for probing differences in the design and performance of nanoparticles. To illustrate this point, consider an example where radiolabeled liposomes were conjugated to Zr^89^ by two different methods [50]. One set of particles were Z^r89^ labeled using a chelator (Zr-SCL), and the other had the isotope conjugated via click-chemistry (Zr-CLL). The authors found that Zr-SCL had greater labeling efficiency, ~6x longer plasma t_1/2_, and greater accumulation in an orthotopic 4T1 breast tumor xenograft. However, fitting tumor accumulation data using the diffusion flux model yielded P values that were not significantly different among the two liposome samples (Figure 10A and B). As such, the labeling techniques employed did not appreciably affect particle permeability in the tumor, and poorer tumor accumulation of can be largely attributed to the particles (or Z^r89^ label) being more quickly eliminated from the bloodstream.

**Figure 10:**
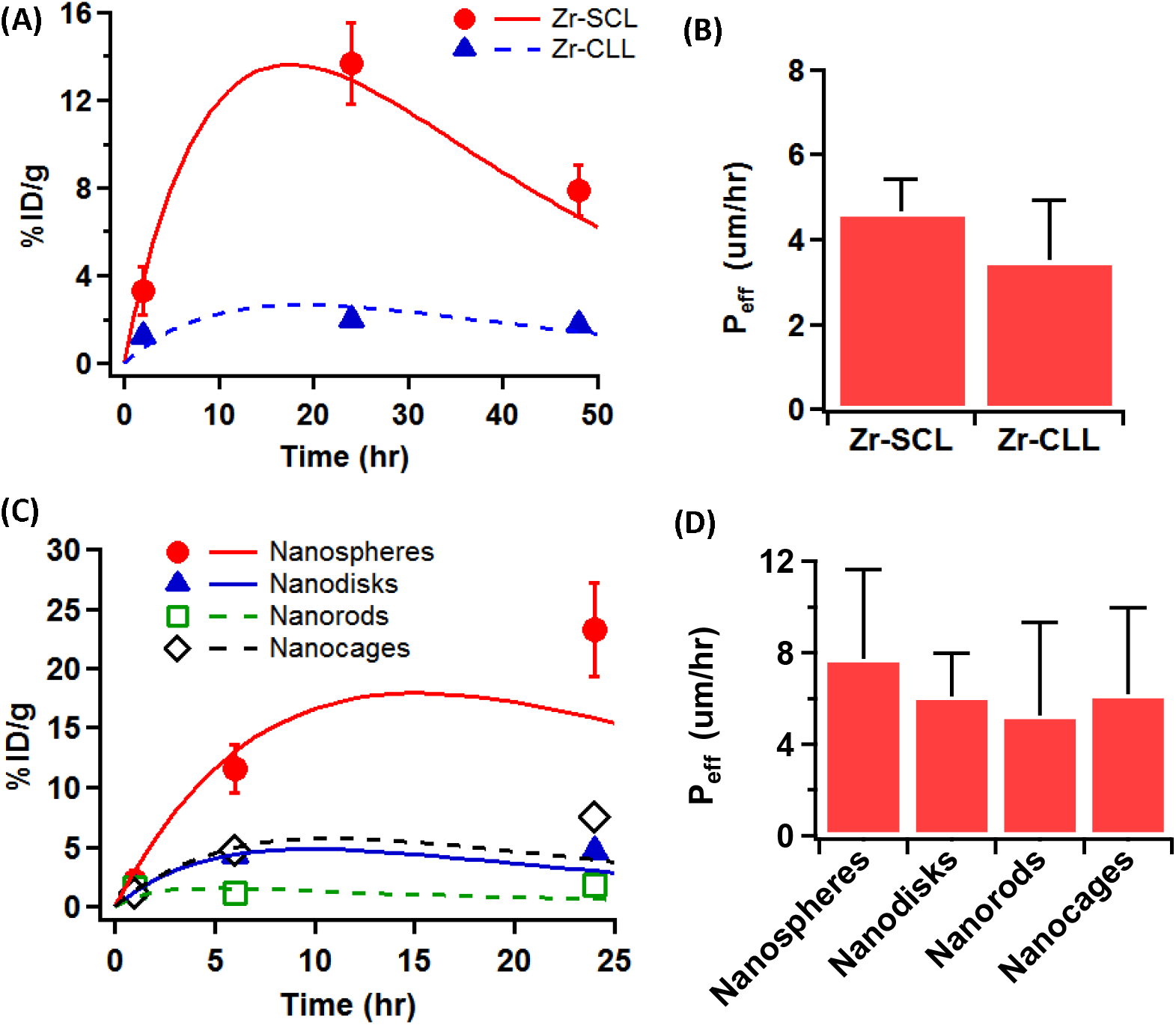
(A) Tumor accumulation of liposomes labeled with Zr^89^ using two different methods, in an orthotopic 4T1 breast cancer model [50], and (B) their fitted P_eff_. (C) Au NP with different shapes tested in EMT6 breast cancer model [54], and (D) their fitted P_eff_. Markers refer to data points gleaned from respective literature sources, while trendlines show fits obtained using diffusion flux model.

In another example, gold nanoparticles (Au NP) synthesized with varying shapes were found to achieve different levels of tumor accumulation in EMT6 breast tumor xenograft [54]. Yet, their *P*_*eff*_ values were also found to have non-significant differences (Figure 10C and D). Therefore, changing the shape of Au NPs did not impact their tumor permeabilities as much as their plasma clearance.

The above scenarios stand in contrast to other systems discussed previously, where the respective plasma half-lives of Dextran chains with various molecular weights did not fully account for their differences in tumor accumulation [32] (Figure 5 and Table 4). It is also distinct from 3HM vs liposomes tested in U87MG GBM model [26], where the particles had similar t_1/2_ but different tumor accumulation owing to different permeabilities (Figure 4).

### 5. Factors leading to poor fits & caveats to the analysis method

In several cases, tumor accumulation data led to poor fits that did not impart a reliable value for *P*_*eff*_ analysis. It is difficult to conclude whether the failure of the fit is due to an inappropriate mathematical model for the system’s physiology (e.g. if convection rather than diffusion predominates transport in those cases), or whether the values reported were themselves problematic. Regardless, some general trends could be observed for factors leading to poorly fitted data, such as when the plasma particle concentrations reported were consistently lower than tumor values [55–57]: a puzzling scenario considering that systemically-administered particles ought to have the highest concentration in the blood pool before being distributed to various organs. The tumor concentration values may also have high variability with no observable trend with respect to time [58,59], or achieved high levels within an hour and appear to plateau despite particle clearance from the bloodstream [60–62], which may indicate lasting retention in the tumors (see examples in Supplementary Information). Some of these cases may have arisen from differences in methods employed for measuring particle concentration in the blood and tumor (e.g. PET imaging analysis vs. chemical extraction and measurement), and also that tumor accumulation may not have excluded contributions from particles in tumor vessels that are distinct from extravasated particles (e.g. if tumor PET signal was reported without correcting for signal from the vasculature, or bulk tumor was processed without first perfusing the animal with saline). However, these factors were similarly not accounted for in the other cases presented earlier, hence being a major caveat of the diffusive flux analysis shown here.

Despite these limitations, in cases where a good fit was achieved, the results demonstrated that using a fast and simple diffusive flux analysis method allowed plasma PK effects to be decoupled from particle permeabilities, enabling a better understanding of factors impacting nanoparticle tumor delivery. Additionally, the method has enabled useful comparisons to be made across different particles in various tumor models conducted by separate research groups, though additional analysis through controlled experiments are needed to confirm the findings.

## Conclusions

In conclusion, herein we have demonstrated a method of analyzing particle tumor transport by fitting to a diffusive flux model. In doing so, we were able to elucidate 3HM transport in several tumor models. The results indicate that 3HM and liposome transport is likely governed by passive diffusion in GBM tumor, considering good fits obtained to the diffusive flux equation. In NDL breast cancer model however, there may be other factors at play (e.g. convective transport) due to the poorer fits. Through this effort, we have determined that 3HM permeability values in GBM, colon, and breast tumor models.

By applying this analysis to other particle data found in literature, we have also demonstrated the first instance of estimating particle permeability using PET data. This stands in contrast to more complex methods employed by other authors, involving cranial window intravital microscopy and compartmental analysis. While the diffusive flux model applied to non-local direct measurements has its drawbacks, the simplicity of this method will allow larger-scale analyses to be made across a growing body of nanoparticle data. As an example, here we have demonstrated meaningful comparisons of permeability values obtained for various particles and tumor models.

## Supplementary Information

The Supplementary Information is available free of charge on XXX at DOI: XXX

## Acknowledgments

Values for particle plasma and tumor concentrations were extracted from several published figures using an open-source software: WebPlotDigitizer, Version 4.1 (https://automeris.io/WebPlotDigitizer), created by Ankit Rohatgi (, Austin, Texas, USA). This work was funded by Tsinghua-UC Berkeley Shenzhen Institute, and the UC Berkeley Chancellor’s Fellowship.

## References

[1] S. Wilhelm, A.J. Tavares, Q. Dai, S. Ohta, J. Audet, H.F. Dvorak, W.C.W. Chan, Analysis of nanoparticle delivery to tumours, Nat. Rev. Mater. 1 (2016) 16014. doi:10.1038/natrevmats.2016.14.

[2] V.J. Venditto, F.C. Szoka, Cancer nanomedicines: So many papers and so few drugs!, Adv. Drug Deliv. Rev. 65 (2013) 80–88. doi:10.1016/j.addr.2012.09.038.

[3] J. Shi, P.W. Kantoff, R. Wooster, O.C. Farokhzad, Cancer nanomedicine: Progress, challenges and opportunities, Nat. Rev. Cancer. 17 (2017) 20–37. doi:10.1038/nrc.2016.108.

[4] L.Y.T. Chou, K. Ming, W.C.W. Chan, Strategies for the intracellular delivery of nanoparticles, Chem. Soc. Rev. 40 (2011) 233–245. doi:10.1039/C0CS00003E.

[5] U. Bulbake, S. Doppalapudi, N. Kommineni, W. Khan, Liposomal formulations in clinical use: An updated review, Pharmaceutics. 9 (2017) 1–33. doi:10.3390/pharmaceutics9020012.

[6] Y. Matsumura, H. Maeda, A new concept for macromolecular therapies in cancer chemotherapy: mechanisms of tumortropic accumulation of proteins and the antitumor agents SMANCS, Cancer Res. 6 (1986) 6392–6397.

[7] H. Maeda, H. Nakamura, J. Fang, The EPR effect for macromolecular drug delivery to solid tumors: Improvement of tumor uptake, lowering of systemic toxicity, and distinct tumor imaging in vivo, Adv. Drug Deliv. Rev. 65 (2013). doi:10.1016/j.addr.2012.10.002.

[8] H. Hashizume, P. Baluk, S. Morikawa, J.W. McLean, G. Thurston, S. Roberge, R.K. Jain, D.M. McDonald, Openings between Defective Endothelial Cells Explain Tumor Vessel Leakiness, Am. J. Pathol. 156 (2000) 1363–1380. doi:10.1016/S0002-9440(10)65006-7.

[9] S.H. Jang, M.G. Wientjes, D. Lu, J.L.S. Au, Drug delivery and transport to solid tumors, Pharm. Res. 20 (2003) 1337–1350. doi:10.1023/A:1025785505977.

[10] R.K. Jain, Vascular and interstitial barriers to delivery of therapeutic agents in tumors, Cancer Metastasis Rev. 9 (1990) 253–266. doi:10.1007/BF00046364.

[11] T. Stylianopoulos, K. Soteriou, D. Fukumura, R.K. Jain, Cationic nanoparticles have superior transvascular flux into solid tumors: Insights from a mathematical model, Ann. Biomed. Eng. 41 (2013) 68–77. doi:10.1007/s10439-012-0630-4.

[12] Y. Boucher, H. Salehil, B. Witwerl, G. Harsh, R. Jain, Interstitial fluid pressure in intracranial tumours in patients and in rodents, Br. Joumal Cancer. 75 (1997) 829–836. doi:10.1038/bjc.1997.148.

[13] M. Stohrer, Y. Boucher, M. Stangassinger, R.K. Jain, Oncotic Pressure in Solid Tumors Is Elevated Oncotic Pressure in Solid Tumors Is Elevated 1, Cancer Res. (2000) 4251–4255.

[14] S.K. Hobbs, W.L. Monsky, F. Yuan, W.G. Roberts, L. Griffith, V.P. Torchilin, R.K. Jain, Regulation of transport pathways in tumor vessels: Role of tumor type and microenvironment, Proc. Natl. Acad. Sci. U. S. A. 95 (1998) 4607–12. doi:10.1073/pnas.95.8.4607.

[15] S.N. Ekdawi, A.S. Mikhail, S. Stapleton, J. Zheng, S. Eetezadi, D.A. Jaffray, C. Allen, Long Circulation and Tumor Accumulation, in: Y.H. Bae, R.J. Mrsny, K. Park (Eds.), Cancer Target. Drug Deliv., Springer Science+Business Media, New York, 2013: pp. 543–571. doi:10.1007/978-1-4614-7876-8.

[16] H. Cabral, Y. Matsumoto, K. Mizuno, Q. Chen, M. Murakami, M. Kimura, Y. Terada, M.R. Kano, K. Miyazono, M. Uesaka, N. Nishiyama, K. Kataoka, Accumulation of sub-100 nm polymeric micelles in poorly permeable tumours depends on size, Nat. Nanotechnol. 6 (2011) 815–823. doi:10.1038/nnano.2011.166.

[17] H. Lee, H. Fonge, B. Hoang, R.M. Reilly, C. Allen, The effects of particle size and molecular targeting on the intratumoral and subcellular distribution of polymeric nanoparticles, Mol. Pharm. 7 (2010) 1195–1208. doi:10.1021/mp100038h.

[18] C. Wong, T. Stylianopoulos, J. Cui, J. Martin, V.P. Chauhan, W. Jiang, Z. Popovic, R.K. Jain, M.G. Bawendi, D. Fukumura, Multistage nanoparticle delivery system for deep penetration into tumor tissue, PNAS. 108 (2011) 2426–2431. doi:10.1073/pnas.1018382108.

[19] H.-J. Li, J.-Z. Du, X.-J. Du, C.-F. Xu, C.-Y. Sun, H.-X. Wang, Z.-T. Cao, X.-Z. Yang, Y.-H. Zhu, S. Nie, J. Wang, Stimuli-responsive clustered nanoparticles for improved tumor penetration and therapeutic efficacy, Proc. Natl. Acad. Sci. 113 (2016) 4164–4169. doi:10.1073/pnas.1522080113.

[20] A. Nagayasu, K. Uchiyama, H. Kiwada, The size of liposomes: a factor which affects their targeting efficiency to tumors and therapeutic activity of liposomal antitumor drugs, Adv. Drug Deliv. Rev. 40 (1999) 75–87.

[21] K. Uchiyama, Y. Yamagiwa, Y. Yamagiwa, T. Nishida, H. Harashima, H. Kiwada, Effects of the size and fluidity of liposomes on their accumulation in tumors!]: A presumption of their interaction with tumors, Int. J. Pharm. 121 (1995) 195–203.

[22] J.N. Israelachvili, D.J. Mitchell, B.W. Ninham, Theory of Self-Assembly of Hydrocarbon Amphiphiles into Micelles and Bilayers, J Chem. Soc. Faraday Trans. 2. 72 (1976) 1525.

[23] C.C. Lee, E.R. Gillies, M.E. Fox, S.J. Guillaudeu, J.M.J. Frechet, E.E. Dy, F.C. Szoka, A single dose of doxorubicin-functionalized bow-tie dendrimer cures mice bearing C-26 colon carcinomas, PNAS. 103 (2006) 16649–16654.https://www.ncbi.nlm.nih.gov/pmc/articles/PMC1636509/bin/pnas_0607705103_index.html#F5

[24] J. Andrew MacKay, M. Chen, J.R. McDaniel, W. Liu, A.J. Simnick, A. Chilkoti, Self-assembling chimeric polypeptide-doxorubicin conjugate nanoparticles that abolish tumours after a single injection, Nat. Mater. 8 (2009) 993–999. doi:10.1038/nmat2569.

[25] M.E. Fox, F.C. Szoka, J.M.J. Fréchet, Soluble Polymer Carrier for the Treatment of Cancer: The Importance of Molecular Architecture, Macromolecules. 42 (2010) 1141–1151. doi:10.1021/ar900035f.Soluble.

[26] J.W. Seo, J. Ang, L.M. Mahakian, S. Tam, B. Fite, E.S. Ingham, J. Beyer, J. Forsayeth, K.S. Bankiewicz, T. Xu, K.W. Ferrara, Self-assembled 20-nm 64Cu-micelles enhance accumulation in rat glioblastoma, J. Control. Release. 220 (2015) 51–60. doi:10.1016/j.jconrel.2015.09.057.

[27] N. Dube, J.Y. Shu, H. Dong, J.W. Seo, E. Ingham, A. Kheirolomoom, P.Y. Chen, J. Forsayeth, K. Bankiewicz, K.W. Ferrara, T. Xu, Evaluation of doxorubicin-loaded 3-helix micelles as nanocarriers, Biomacromolecules. 14 (2013) 3697–3705. doi:10.1021/bm4010518.

[28] H. Dong, N. Dube, J.Y. Shu, K.W. Ferrara, T. Xu, Long-Circulating 15 nm Micelles Based on Amphiphilic 3-Helix Peptide-PEG Conjugates, ACS Nano. 6 (2012) 5320–5329. doi:10.1021/nn391142r.

[29] H. Dong, J.Y. Shu, N. Dube, Y. Ma, M. V. Tirrell, K.H. Downing, T. Xu, 3-Helix micelles stabilized by polymer springs, J. Am. Chem. Soc. 134 (2012) 11807–11814. doi:10.1021/ja3048128.

[30] J.W. Seo, H. Zhang, D.L. Kukis, C.F. Meares, K.W. Ferrara, A Novel Method to Label Preformed Liposomes with (CU)-C-64 for Positron Emission Tomography (PET) Imaging, Bioconjug. Chem. 19 (2008) 2577–2584. doi:10.1021/bc8002937.

[31] N. Anatoly, P. Vladimir, Polyethylene glycol-diacyllipid micelles demonstrate increased accumulation in subcutaneous tumors in mice, Pharm. Res. 19 (2002) 1424–1429.

[32] M.R. Dreher, W. Liu, C.R. Michelich, M.W. Dewhirst, F. Yuan, A. Chilkoti, Tumor vascular permeability, accumulation, and penetration of macromolecular drug carriers, J. Natl. Cancer Inst. 98 (2006) 335–344. doi:10.1093/jnci/djj070.

[33] F. Yuan, M. Leunig, S.K. Huang, D.A. Berk, D. Papahadjopoulos, R.K. Jam, Microvascular Permeability and Interstitial Penetration of Sterically Stabilized (Stealth) Liposomes in a Human Tumor Xenograft ’, Cancer Res. 54 (1994) 3352–3357.

[34] R.K. Jain, Transport of molecules across tumor vasculature, Cancer Metastasis Rev. 6 (1987).559–593. doi:10.1007/BF00047468.

[35] O. Chen, V.P. Chauhan, T. Stylianopoulos, J.D. Martin, Z. Popovic, W.S. Kamoun, M.G. Bawendi, D. Fukumura, R.K. Jain, Normalization of tumour blood vessels improves the delivery of nanomedicines in a size-dependent manner, Nat. Nanotechnol. 7 (2012) 383–388. doi:10.1038/nnano.2012.45.

[36] N.Z. Wu, D. Da, T.L. Rudoll, S. Liposomes, D. Needham, A.R. Whorton, M.W. Dewhirst, Increased Microvascular Permeability Contributes to Preferential Accumulation of Stealth Liposomes in Tumor Tissue Increased Microvascular Permeability Contributes to Preferential Accumulation of, Cancer Res. 53 (1993) 3765–3770.

[37] N. Dube, J.W. Seo, H. Dong, J.Y. Shu, R. Lund, L.M. Mahakian, K.W. Ferrara, T. Xu, Effect of Alkyl Length of Peptide − Polymer Amphiphile on Cargo Encapsulation Stability and Pharmacokinetics of 3 IZ Helix Micelles, Biomacromolecules. 15 (2014) 2963–2970.

[38] Y. Kuang, J.D. Nagy, S.E. Eikenberry, Introduction to Mathematical Oncology, CRC Press, Boca Raton, FL, 2016.

[39] D. Fukumura, R.K. Jain, Tumor microenvironment abnormalities: Causes, consequences, and strategies to normalize, J. Cell. Biochem. 101 (2007) 937–949. doi:10.1002/jcb.21187.

[40] C.-H. Heldin, K. Rubin, K. Pietras, A. Ostman, High interstitial fluid pressure - an obstacle in cancer therapy., Nat. Rev. Cancer. 4 (2004) 806–13. doi:10.1038/nrc1456.

[41] D.J. Wolak, R.G. Thorne, Diffusion of Macromolecules in the Brain: Implications for Drug Delivery, Mol. Pharm. 10 (2013) 1492–1504. doi:10.1021/mp300495e.

[42] R.T. Tong, Y. Boucher, S. V Kozin, F. Winkler, D.J. Hicklin, R.K. Jain, Vascular Normalization by Vascular Endothelial Growth Factor Receptor 2 Blockade Induces a Pressure Gradient Across the Vasculature and Improves Drug Penetration in Tumors, Cancer Res. 64 (2004) 3731–3736.

[43] F. Yuan, H.A. Salehi, Y. Boucher, R.K. Jain, U.S. Vasthare, R.F. Tuma, Vascular Permeability and Microcirculation of Gliomas and Mammary Carcinomas Transplanted in Rat and Mouse Cranial Windows, Cancer Res. 54 (1994) 4564–4568.

[44] H. Dong, N. Dube, J.Y. Shu, J.W. Seo, L.M. Mahakian, K.W. Ferrara, T. Xu, Long-circulating 15 nm micelles based on amphiphilic 3-helix peptide-peg conjugates, ACS Nano. 6 (2012) 5320–5329. doi:10.1021/nn301142r.

[45] A. Hawe, W.L. Hulse, W. Jiskoot, R.T. Forbes, Taylor dispersion analysis compared to dynamic light scattering for the size analysis of therapeutic peptides and proteins and their aggregates, Pharm. Res. 28 (2011) 2302–2310. doi:10.1007/s11095-011-0460-3.

[46] H. Maeda, Toward a full understanding of the EPR effect in primary and metastatic tumors as well as issues related to its heterogeneity, Adv. Drug Deliv. Rev. 91 (2015) 3–6. doi:10.1016/j.addr.2015.01.002.

[47] N.Z. Wu, B. Klitzman, G. Rosner, D. Needham, M.W. Dewhirst, Measurement of Material Extravasation in Microvascular Networks Using Fluorescence Video-Microscopy, Microvasc. Res. 46 (1993) 231–253. doi:10.1006/mvre.1993.1049.

[48] A. Pathak, P. Kumar, K. Chuttani, S. Jain, A.K. Mishra, S.P. Vyas, K.C. Gupta, Gene Expression, Biodistribution, and Pharmacoscintigraphic Evaluation of Chondroitin Sulfate-PEI Nanoconstructs Mediated Tumor Gene Therapy, ACS Nano. 3 (2009) 1493–1505.

[49] A.W. Wong, E. Ormsby, H. Zhang, J.W. Seo, L.M. Mahakian, C.F. Caskey, K.W. Ferrara, A comparison of image contrast with 64Cu-labeled long circulating liposomes and 18F-FDG in a murine model of mammary carcinoma, Am. J. Nucl. Med. Mol. Imaging. 3 (2013) 32–43. doi:10.2522/ptj.20100142.

[50] C. Pérez-Medina, D. Abdel-Atti, Y. Zhang, V.A. Longo, C.P. Irwin, T. Binderup, J. Ruiz-Cabello, Z.A. Fayad, J.S. Lewis, W.J.M. Mulder, T. Reiner, A Modular Labeling Strategy for In Vivo PET and Near-Infrared Fluorescence Imaging of Nanoparticle Tumor Targeting, J. Nucl. Med. 55 (2014).1706–1711. doi:10.2967/jnumed.114.141861.

[51] Y. Miyajima, H. Nakamura, Y. Kuwata, J. Lee, S. Masunaga, K. Ono, K. Maruyama, Transferrin-Loaded nido-Carborane Liposomes⍰: Tumor-Targeting Boron Delivery System for Neutron Capture Therapy, Bioconjug. Chem. 17 (2006) 1314–1320. doi:10.1021/bc060064k.

[52] Y.-Y. Lin, C.-H. Chang, J.-J. Li, M.G. Stabin, Y.-J. Chang, L.-C. Chen, M.-H. Lin, Y.-L. Tseng, W.-J. Lin, T.-W. Lee, G. Ting, C.A. Chang, F.-D. Chen, H.-E. Wang, Pharmacokinetics and Dosimetry of ^111^ In/ ^188^ Re-Labeled PEGylated Liposomal Drugs in Two Colon Carcinoma-Bearing Mouse Models, Cancer Biother. Radiopharm. 26 (2011) 373–380. doi:10.1089/cbr.2010.0906.

[53] B.L. Coomberl, P.A. Stewart, K. Hayakawa, C.L. Farrell, R.F. Del Maestros, Quantitative morphology of human glioblastoma multiforme microvessels: structural basis of blood-brain barrier defect, J. Neurooncol. 5 (1987) 299–307. doi:10.1007/BF00148386.

[54] K.C.L. Black, Y. Wang, H.P. Luehmann, X. Cai, W. Xing, B. Pang, Y. Zhao, C.S. Cutler, L. V. Wang, Y. Liu, Y. Xia, Radioactive198Au-doped nanostructures with different shapes for in vivo analyses of their biodistribution, tumor uptake, and intratumoral distribution, ACS Nano. 8 (2014).4385–4394. doi:10.1021/nn406258m.

[55] L.M. Negi, S. Talegaonkar, M. Jaggi, A. Kamra, R. Verma, S. Dobhal, V. Kumar, Surface engineered nanostructured lipid carriers for targeting MDR tumor⍰: Part II. In vivo biodistribution, pharmacodynamic and hematological toxicity studies, Colloids Surfaces B Biointerfaces. 123 (2014) 610–615. doi:10.1016/j.colsurfb.2014.09.061.

[56] Y. Chen, W. Zhang, Y. Huang, F. Gao, X. Sha, X. Fang, Pluronic-based functional polymeric mixed micelles for co-delivery of doxorubicin and paclitaxel to multidrug resistant tumor, Int. J. Pharm. 488 (2015) 44–58. doi:10.1016/j.ijpharm.2015.04.048.

[57] J. Xu, F. Gattacceca, M. Amiji, Biodistribution and pharmacokinetics of EGFR-targeted thiolated gelatin nanoparticles following systemic administration in pancreatic tumor-bearing mice, Mol. Pharm. 10 (2013) 2031–2044. doi:10.1021/mp400054e.

[58] C.M. Dawidczyk, L.M. Russell, M. Hultz, P.C. Searson, Tumor accumulation of liposomal doxorubicin in three murine models: Optimizing delivery efficiency, Nanomedicine Nanotechnology, Biol. Med. 13 (2017) 1637–1644. doi:10.1016/j.nano.2017.02.008.

[59] C. Tsai, C. Chang, L. Chen, Y.-J. Chang, K.-L. Lan, Y. Wu, C.-W. Hsu, I. Liu, C.-L. Ho, W. Lee, H. Ni, T.-J. Chang, G. Ting, T.-W. Lee, Biodistribution and pharmacokinetics of 188 Re-liposomes and their comparative therapeutic efficacy with 5-fluorouracil in C26 colonic peritoneal carcinomatosis mice, Int. J. Nanomedicine. 6 (2011) 2607–2619.

[60] K. Cheng, S.R. Kothapalli, H. Liu, A.L. Koh, J. V. Jokerst, H. Jiang, M. Yang, J. Li, J. Levi, J.C. Wu, S.S. Gambhir, Z. Cheng, Construction and validation of nano gold tripods for molecular imaging of living subjects, J. Am. Chem. Soc. 136 (2014) 3560–3571. doi:10.1021/ja412001e.

[61] H. Hong, K. Yang, Y. Zhang, J.W. Engle, L. Feng, Y. Yang, T.R. Nayak, S. Goel, J. Bean, C.P. Theuer, T.E. Barnhart, Z. Liu, W. Cai, In Vivo Targeting and Imaging of Tumor Vasculature with Radiolabeled, Antibody-Conjugated Nanographene, ACS Nano. 6 (2012) 2361–2370. doi:10.1021/nn204625e.

[62] S. Shi, K. Yang, H. Hong, H.F. Valdovinos, T.R. Nayak, Y. Zhang, C.P. Theuer, T.E. Barnhart, Z. Liu, W. Cai, Tumor vasculature targeting and imaging in living mice with reduced graphene oxide, Biomaterials. 34 (2013) 3002–3009. doi:10.1016/j.biomaterials.2013.01.047.

