## Supplemental Information for "Diffusive Flux Analysis of Tumor Vascular Permeability for 3-Helix-Micelles in Comparison to Other Nanoparticles"

Supplementary Information

##### 1. Effect of active targeting

Diffusive flux modeling can also be used to compare the effect of active-targeting surface ligands on particle tumor permeability. Various strategies taking advantage of tumor-specific overexpression of receptors have been published in literature. Ligands include a monoclonal antibody for binding to CD105 tumor endothelial marker for angiogenesis (TRC105)[1,2], epidermal growth factor (EGF) for facilitating nuclear translocation in epithelial cancers [3,4], folic acid (FA) for binding to high-affinity folate receptors [5,6]; and transferrin for binding to transferrin receptors [7]. In all these cases, ligand-receptor binding is expected to enhance particle and cargo exposure to tumor cells compared to their untargeted counterparts. Despite the overall similarity of targeting approaches, some differences could be seen in the permeability of particles decorated with these various ligands, with FA targeting having the most pronounced effect in driving tumor accumulation.

EGF targeting for PEG-polycaprolactone block copolymer micelles (BCPM) appear to modestly increase in $P_{eff}$ in MDA breast cancer models [3], while Tf targeting for Nido-carborane liposomes (NidoC Lipo) in a colon cancer model [7] did not have an appreciable impact on $P_{eff}$ compared to their nontargeted counterparts (Figure S1). In the EGF targeting experiment, two different breast cancer models were employed: MCF7 which expresses ~100x less EGF receptors (10^4^ EGFR/cell) than MDA-MB-468 (10^6^ EGFR/cell) serves as a control for the targeting strategy [3]. As expected, tumor accumulation was higher for EGF targeted BCPM in MDA-MB-468 tumor, while no appreciable difference was observed in MCF7 tumor (Figure S1A and B). Interestingly, $P_{eff}$ values for these BCPM particles are noticeably lower than typical for most breast cancer models (see Table S1), and the reasons for this are not immediately apparent.

**(B)**

**(A)**

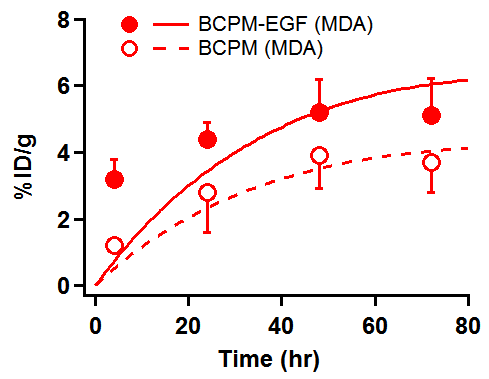

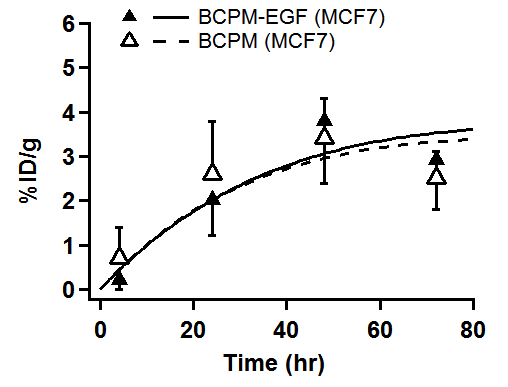

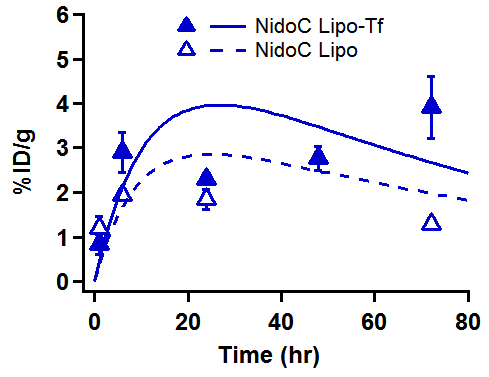

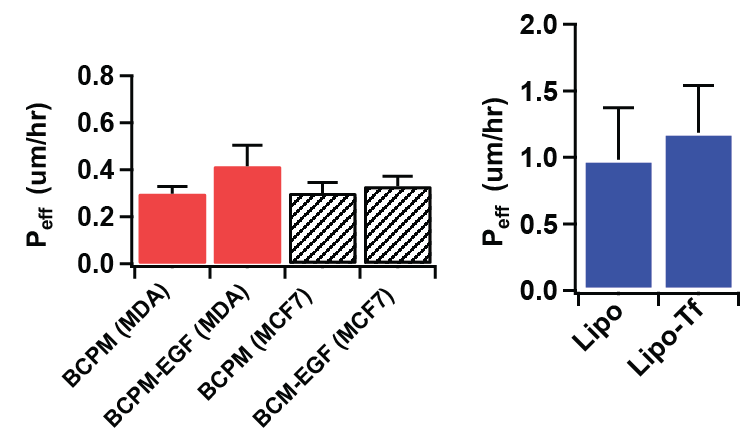

**(E)**

**(D)**

**(C)**

Figure S1: Tumor accumulation plots for PEG-PCL BCPM with and without EGF targeting dosed to (A) MDA-MB-468 and (B) MCF-7 subcutaneous breast tumor models, as reported by Lee et al [3]. (C) Tumor accumulation of Nido-carborane liposomes with and without Transferrin (Tf) ligand in C26 colon tumor model, reported by Miyajima et al [7]. Data points were extracted from the respective articles, and trendlines represent Peff fits using the diffusion flux model; see Table S1 for R^2^ of fits. (D) $P_{eff}$values for both breast tumor models and (E) for colon tumor model.

Targeting using CD105 tumor endothelial marker (TRC105) appear to enhance $P_{eff}$ of reduced graphene oxide (RGO) and poly(amidoamine)-poly(L- lactide)-b-poly(ethylene glycol) (PAMAM-PLA-PEG) block copolymer particles in 4T1 subcutaneous breast tumor models (Figure S2). However, the fits for non-targeted series appear to be poor and hence their $P_{eff}$ values are non-trustworthy.

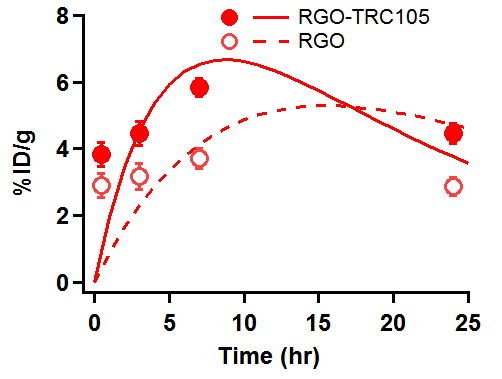

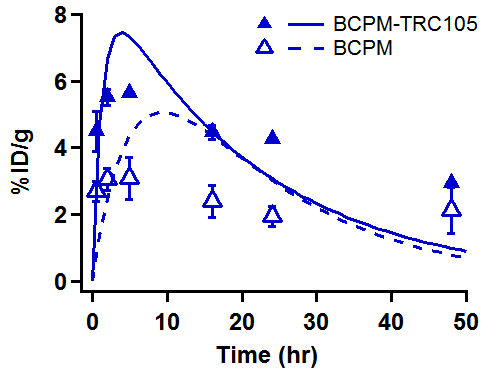

**(B)**

**(A)**

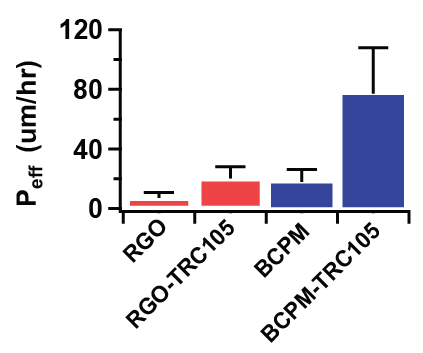

**(C)**

Figure S2: Tumor accumulation plots for (A) RGO particles [1] and (B) PAMAM-PLA-PEG block copolymer micelles [2] with and without TRC105 targeting dosed 4T1 subcutaneous breast tumor model. Data points were extracted from the respective articles, and trendlines represent $P_{eff}$ fits using the diffusion flux model; see Table S1 for R^2^ of fits. (C) Peff values for fits in (A) and (B) show statistically significant difference between targeted and untargeted particles.

Unlike the previous cases, particles with folic acid (FA) targeting ligand appear to be capable of exceeding accumulation by simple diffusive transport. This is observed in two separate cases with PEG-PCL block copolymer micelles [5] and PLGA nanoparticles [6] dosed to 4T1 and MDA-MB-231 breast tumor models. As shown in Figure S3 A & B, FA-targeted particles accumulated in the tumor beyond the best fit trendlines modeled by diffusion flux. This can be seen to a lesser extent in NGR (CD31) peptide-targeted liposomes dosed to non-small cell lung cancer model[8]. As shown in Figure S3 C & D, liposomes doped with NGR-PEG lipids had higher tumor accumulation and effective permeability compared to controls of untargeted liposomes and liposomes doped with mismatched ARA peptide. However, the liposomes with longer PEG 3400 chain presenting NGR ligand appear to accumulate in the tumor slightly beyond diffusive flux.

**(B)**

**(A)**

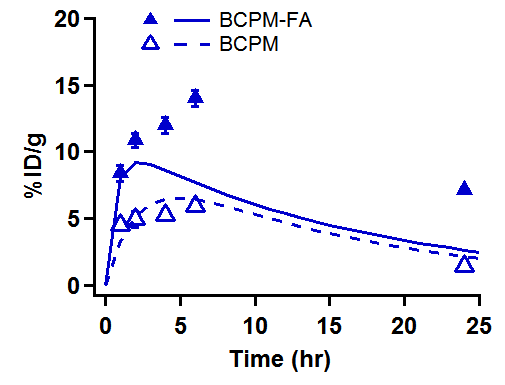

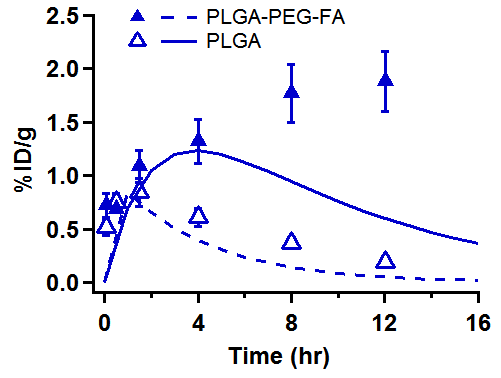

**(D)**

**(C)**

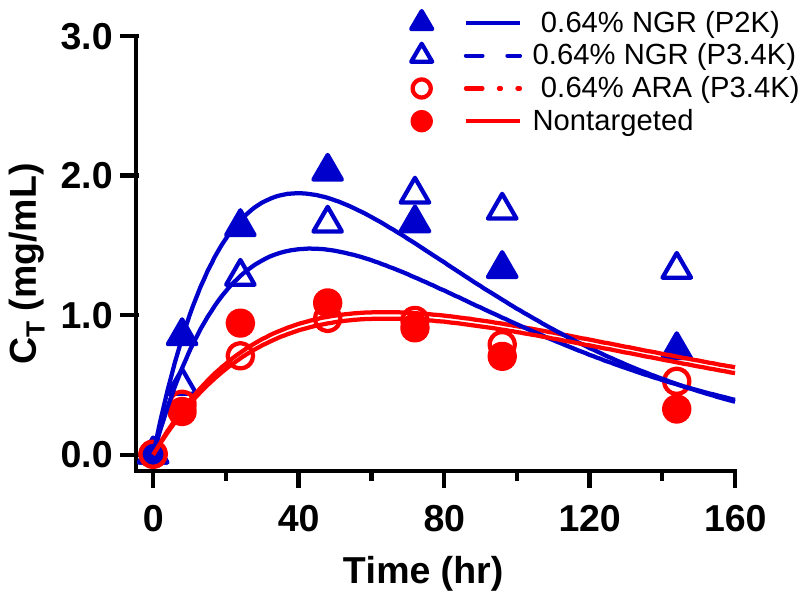

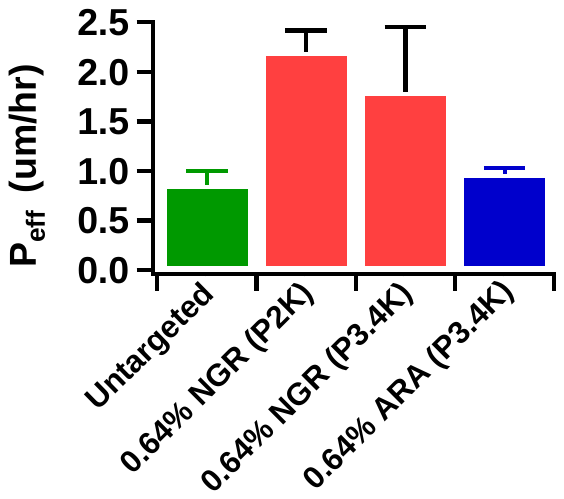

Figure S3: Tumor accumulation plots for (A) PEG-PCL block copolymer micelles dosed to 4T1 subcutaneous breast tumor model [5], and (B) PLGA nanoparticles dosed to MDA-MB-231 breast tumor model[6], prepared with and without folic acid (FA) targeting ligand. (C)&(D) Tumor accumulation and $P_{eff}$ of PEGylated liposomes with NGR or mismatched ARA peptide dosed to H520 non-small cell lung cancer model[8]. Data points were extracted from the respective articles, and trendlines represent $P_{eff}$ fits using the diffusion flux model; see Table S1 for R^2^ of fits. Arrows point to targeted particle series rising above best trendline fit possible.

The apparent differences in tumor transport between EGF, TRC105, Tf, NGR and FA-targeted particles is surprising. Inducing particle endocytosis via ligand-receptor coupling is expected to drive transport by creating a perpetual “sink” condition in the extravascular space that increases the concentration gradient between vascular vs. interstitial fluid concentration. When vascular particle concentration drops from elimination pathways, the concentration gradient typically reverses, with higher concentration in the extravascular space driving diffusion of particles back into the vessels. However, endocytosed particles are not susceptible to being washed out by this method, and therefore tumor accumulation should remain high. As such tumor accumulation curves presented in Figure S3 for targeted vs. nontargeted particles is the expected scenario for active-targeted endocytosed particles, suggesting that FA-targeting strategy may be superior over other ligands. Further study in a controlled environment with matching particles and tumor models would need to be performed to ascertain this conclusion.

Note that the above analysis on active-targeting has only dealt with tumor accumulation; there is also evidence to suggest that targeting ligands can adversely impact tumor penetration, owing to the binding-site barrier [4,9]. That is, the fact that ligand-conjugated particles will adhere to tumor cells encountered soon after extravasation actually prevents the particles from diffusing further into the tissue, unless a saturating dose is administered to occupy all binding sites [10]. Insufficient tumor penetration has been attributed to disappointing therapeutic benefit of the FDA-approved Doxil and Abraxane which are ≤100 nm[11]. As such systems were developed to enable nanoparticle size to be tuned from the classical ~100 nm design to release smaller particles and thus improve tumor penetration [12], and a recent example with acid-labile particles demonstrated greater efficacy in a poorly permeable pancreatic cancer model [13]. As such, the use of targeting ligands should be explored with caution, paying to mind the cost-benefit ratio of increasing tumor accumulation (if any) at the cost of limiting tumor penetration.

##### 2. Fitting with and without using standard deviation of data points

Ideally, $P_{eff}$ fitting is done by taking into account the standard deviation (SD) of $C_{T}$values each time-point. This could work in some cases, however if a point has a much smaller SD than the others, the Python algorithm (via SciPy “curve_fit” function) performs poorly. Consider the following cases of 3HM and liposome accumulation data in U87MG GBM model, reproduced below from the main text (Figure S4). If the mean values of all points were assumed to be nominal values, the algorithm will attempt a fit without bias to any point, yielding P_eff_ for 3HM = 0.832 ± 0.065, and for Liposome = 0.389 ± 0.010.

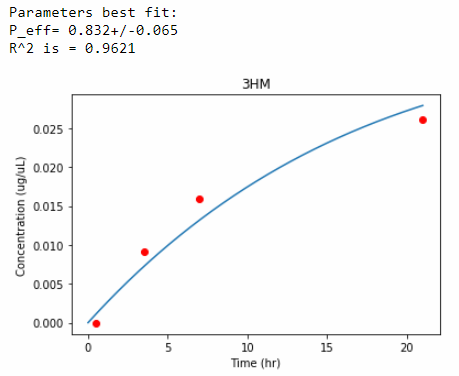

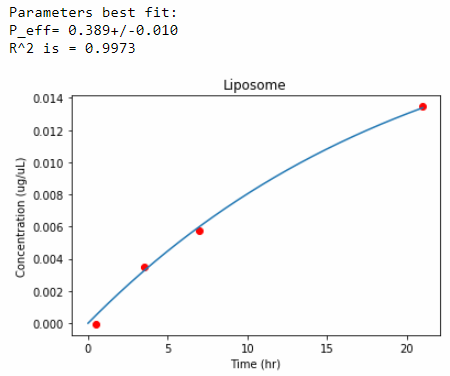

Figure S4: 3HM and liposome tumor accumulation fits (trend-lines), without considering SD of individual data points (markers)

However, including the original SD for the t = 0.5 hr time-point (<0.001 ug/uL) leads to a poor fit. This yields P_eff_ for 3HM = 0.230 ± 0.234, and for Liposome = 0.196 ± 0.130.

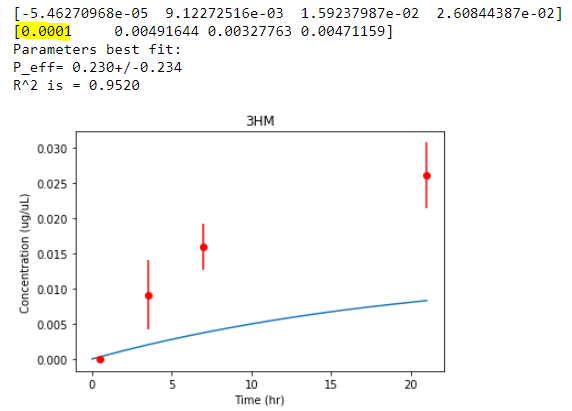

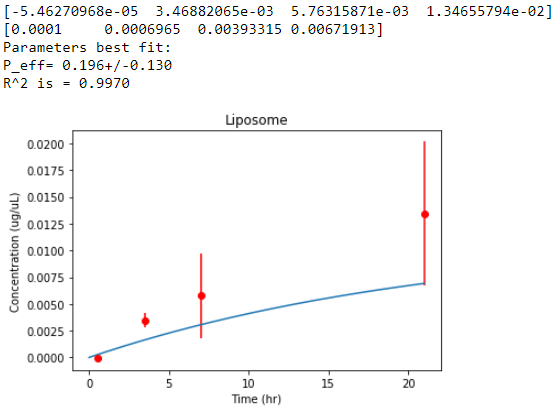

Figure S5: 3HM and liposome tumor accumulation fits (trend-lines), considering SD of individual data points (markers with error bars)

If the SD at t = 0.5 hr is artificially set to 0.001 ug/mL, the fit is restored, yielding P_eff_ for 3HM = 0.838 ± 0.108, and for Liposome = 0.405 ± 0.026

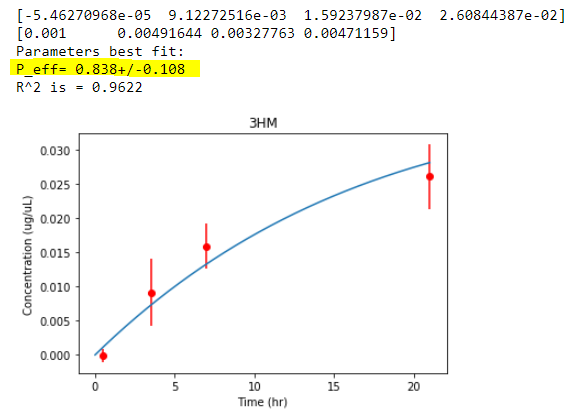

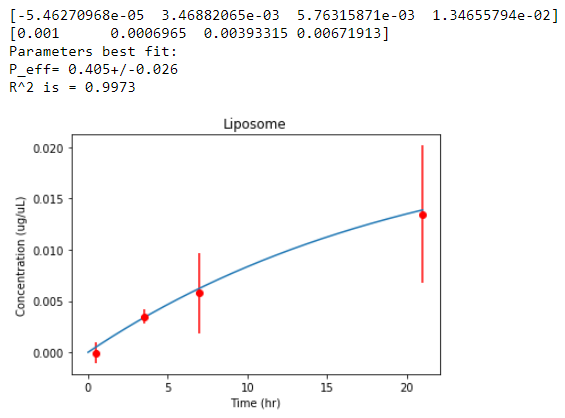

Figure S6: 3HM and liposome tumor accumulation fits (trend-lines), considering SD of individual data points (markers with error bars). SD at t = 0.5 hr was artificially set to 0.001 ug/uL.

The difference in P_eff_ between fits shown in Figure S4 and Figure S6 is < 5%, but the fit error values differ by 40% and 62% for 3HM and Liposome series, respectively. This is expected, since the error of the individual data points contribute to the overall error of $P_{eff}$. While this is the ideal scenario, many other data sets compared in this paper have non-uniform SD values that lead to poor fits. Artificially setting SD values that restore the fit for each case would be disingenuous and impractical. As such, we have elected to uniformly fit all data sets by assuming the mean $C_{T}$ values as nominal values, with the caveat that $P_{eff}$error values produced from the fits are underestimated.

##### 3. Model Diagnostics

$P_{eff}$ is estimated by minimizing the least square error of Equation 9 in the main text. As in standard regression analysis, here we assume the noise are independently identically distributed Gaussian. That is, $C_{T}\left( t \right)=f\left( t, P_{eff} \right)+\varepsilon$, where $\epsilon\sim N\left( 0,\sigma^{2} \right)$ and according to Equation 9:

$$f\left( t, P_{eff} \right)=\frac{AP_{eff}C_{0}/V_{T}}{K-AP_{eff}/V_{T}}\left( e^{-\frac{AP_{eff}}{V_{T}}t}-e^{-Kt} \right)$$

Our least square estimator $\hat{P}_{eff}$ will be Gaussian distributed where the estimated variance is given by delta method. To compare $\hat{P}_{eff}$ between fits, we applied two-sample t-test. However, a few factors (see Section 5 in the main text) may introduce bias in our model, which makes the t-test invalid. We thus used residual plots to help identify the unexplained variation. In Figure S7, we plot the time versus the residuals for BCP micelles in a pancreatic cancer model[11]. There is a significant effect of time in the residuals with a significance level of 0.000037. This is an example where our model fails to explain the data. As a result, we cannot trust the p-value given by the t-test for differentiating $\hat{P}_{eff}$ among the fits. On the other hand, Figure S8 shows an example from 3HM and Liposomes in a GBM tumor model[14] where the residuals do not exhibit a time trend and are randomly scattered around zero. This indicates that we made a valid model assumption as stated above. Thus, the estimated $\hat{P}_{eff}$ and the t-test comparing 3HM and liposome might be trustworthy.

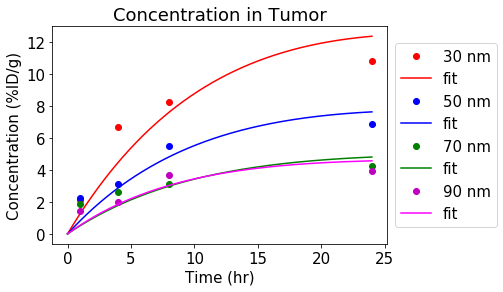

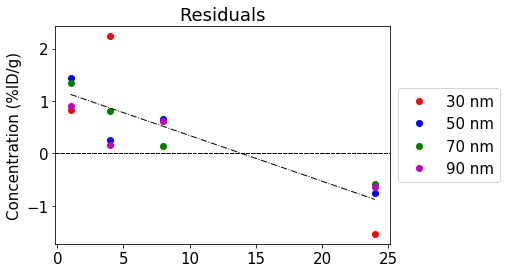

Figure S7: Tumor accumulation profiles and residuals of the fit for BCP micelles in subcutaneous BxPC3 pancreatic cancer.

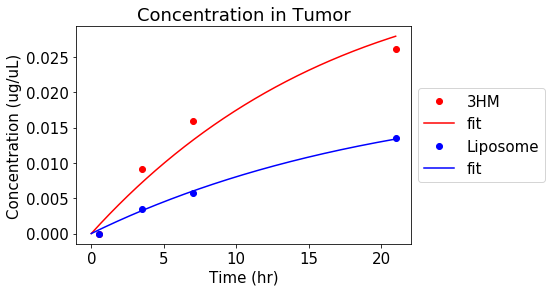

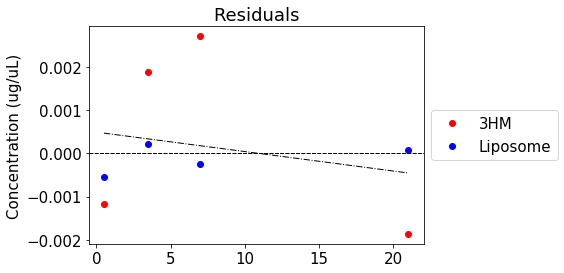

Figure S8: Tumor accumulation profiles and residuals of the fit for 3HM and Liposomes in orthotopic U87MG GBM.

##### 4. Compiled permeability and fit parameter values

Table S1: Permeability and fit parameter values. (OD: original data, SubQ: subcutaneous, BSA: bovine serum albumin, BCP: block copolymer, FA: folic acid, PEG: polyethyleneglycol, SD: standard deviation, K: plasma elimination time constant, PK: pharmacokinetics, * based on reported value from source article, ** poor fit)

| **Ref** | **Tumor** | **Implant** | **Cell Line** | **Particle** | **Size (nm)** | **Active/**  **Passive** | **Ligand Type** | **P_eff_ (um/hr)** | **P_eff_ SD (um/hr)** | **P_eff_ (cm/s)** | **P_eff_ SD (cm/s)** | **K (hr^-1^)** | **C_0_** | **Units** | **PK Fit R^2^** | **# Time Pts.** |
| --- | --- | --- | --- | --- | --- | --- | --- | --- | --- | --- | --- | --- | --- | --- | --- | --- |
| [15] | Breast | SubQ | MCAIV | Albumin-TRITC BSA | 7 | Passive | None | 24.52* | 4.46* | 6.81E-07* | 1.24E-07* | N/A | N/A | N/A | N/A | N/A |
| [16] | Breast | SubQ | R3230AC | PEG Liposome | 100 | Passive | None | 12.3* | 2.81* | 3.42E-07* | 7.80E-08* | N/A | N/A | N/A | N/A | N/A |
| [16] | Breast | SubQ | R3230AC | Liposome | 100 | Passive | None | 6.3* | 1.37* | 1.75E-07* | 3.80E-08* | N/A | N/A | N/A | N/A | N/A |
| [1] | Breast | SubQ | 4T1 | ^66^Ga-NOTA-RGO-TRC105 (targeting) | 27 ± 0.9 | Active | TRC-105 | 20.3 | 7.89 | 5.64E-07 | 2.19E-07 | 0.0570 | 10.952 | %ID/g | 0.7601 | 4 |
| [1] | Breast | SubQ | 4T1 | ^66^Ga-NOTA-RGO (nontargeting) | 27 ± 0.9 | Passive | None | 7.298** | 3.553 | 2.03E-07** | 9.87E-08 | 0.0570 | 12.777 | %ID/g | 0.8922 | 4 |
| [2] | Breast | SubQ | 4T1 | PAMAM-PLA-PEG BCP micelles (TRC105 targeting) | 37 | Active | TRC-105 | 75.36 | 32.29 | 2.09E-06 | 8.97E-07 | 0.0470 | 8.9629 | %ID/g | 0.8978 | 6 |
| [2] | Breast | SubQ | 4T1 | PAMAM-PLA-PEG BCP micelles (no targeting) | 37 | Passive | None | 16.12** | 7.56 | 4.48E-07** | 2.10E-07 | 0.0580 | 8.6991 | %ID/g | 0.9097 | 6 |
| [5] | Breast | SubQ | 4T1 | PEG-PCL micelle (control, no FA) | ~70 | Passive | None | 49.24 | 10.92 | 1.37E-06 | 3.03E-07 | 0.0640 | 8.859 | %ID/g | 0.9104 | 5 |
| [5] | Breast | SubQ | 4T1 | PEG-PCL with FA and carbamate Dox linker | 86.6 | Active | FA | 213.3** | 475.3 | 5.93E-06** | 1.32E-05 | 0.0540 | 7.3691 | %ID/g | 0.8652 | 5 |
| [5] | Breast | SubQ | 4T1 | PEG-PCL with FA and hydrazone Dox linker | 70.9 | Active | FA | 156.1** | 192.4 | 4.34E-06** | 5.34E-06 | 0.0590 | 10.533 | %ID/g | 0.9119 | 5 |
| [6] | Breast | SubQ | MDA-MB-231 | PLGA (control no FA or PEG) | 217-289 | Passive | None | 691.9 | 357.1 | 1.92E-05 | 9.92E-06 | 0.2530 | 1.0419 | ug/g | 0.8312 | 6 |
| [6] | Breast | SubQ | MDA-MB-231 | PLGA (FA + PEG) | 217-289 | Active | FA+PEG | 45.83** | 42.33 | 1.27E-06** | 1.18E-06 | 0.1260 | 2.0139 | ug/g | 0.8479 | 6 |
| [3] | Breast | SubQ | MDA-MB-468 | PEG-PCL micelle no targeting | 57 ± 8 | Passive | None | 0.299 | 0.030 | 8.31E-09 | 8.33E-10 | 0.0230 | 45.956 | %ID/g | 0.9017 | 4 |
| [3] | Breast | SubQ | MDA-MB-468 | PEG-PCL micelle with targeting (T-BCM) | 61 ± 1 | Active | T-BCM | 0.466 | 0.095 | 1.29E-08 | 2.64E-09 | 0.0240 | 45.762 | %ID/g | 0.9907 | 4 |
| [3] | Breast | SubQ | MCF-7 | PEG-PCL micelle no targeting | 57 ± 8 | Passive | None | 0.301 | 0.045 | 8.36E-09 | 1.25E-09 | 0.0260 | 39.095 | %ID/g | 0.8904 | 4 |
| [3] | Breast | SubQ | MCF-7 | PEG-PCL micelle with targeting (T-BCM) | 61 ± 1 | Active | T-BCM | 0.330 | 0.042 | 9.17E-09 | 1.17E-09 | 0.0220 | 34.291 | %ID/g | 0.9583 | 4 |
| [17] | Breast | SubQ | EMT6 | Au Nanospheres | 56.8 | Passive | None | 7.745 | 3.888 | 2.15E-07 | 1.08E-07 | 0.0560 | 41.791 | %ID/g | 0.959 | 3 |
| [17] | Breast | SubQ | EMT6 | Au Nanodisks | 92 x 7 | Passive | None | 6.096 | 1.882 | 1.69E-07 | 5.23E-08 | 0.1580 | 22.712 | %ID/g | 0.9933 | 3 |
| [17] | Breast | SubQ | EMT6 | Au Nanorods | 39.2 L x 9.2 diam | Passive | None | 5.264 | 4.074 | 1.46E-07 | 1.13E-07 | 0.5040 | 19.096 | %ID/g | 0.9987 | 3 |
| [17] | Breast | SubQ | EMT6 | Au Nanocages | 49.6 L x 4.5 thick | Passive | None | 6.185 | 3.776 | 1.72E-07 | 1.05E-07 | 0.1280 | 23.332 | %ID/g | 0.9667 | 3 |
| [18] | Breast | SubQ | EAT | Branched PEI with chondroitin sulfate | 114 | Passive | None | 43.30 | 7.83 | 1.20E-06 | 2.18E-07 | 0.0460 | 0.372 | ug/g | 0.9687 | 4 |
| [19] | Breast | Ortho-topic, breast | MET1 | PEG Liposome | 100 | Passive | None | 2.567 | 0.976 | 7.13E-08 | 2.71E-08 | 0.0460 | 56.05582 | mg/mL | 0.9988 | 3 |
| [20] | Breast | Ortho-topic, breast | NDL | 3HM-C16 | 17 | Passive | None | 1.028** | 0.395 | 2.86E-08** | 1.10E-08 | 0.0440 | 0.2854 | mg/mL | 0.9579 | 5 |
| [20] | Breast | Ortho-topic, breast | NDL | 3HM-C18 | 17.6 | Passive | None | 0.654** | 0.206 | 1.82E-08** | 5.72E-09 | 0.0250 | 0.3294 | mg/mL | 0.9825 | 5 |
| [21] | Breast | Ortho-topic, breast | 4T1 | Zr^89^ Liposome CLL (click chemistry) | 103.4 ± 5.1 | Passive | None | 3.498 | 1.441 | 9.72E-08 | 4.00E-08 | 0.0630 | 8.6707 | %ID/g | 0.8476 | 3 |
| [21] | Breast | Ortho-topic, breast | 4T1 | Zr^89^ Liposome SCL (chelator chemistry) | 103.4 ± 5.1 | Passive | None | 4.638 | 0.767 | 1.29E-07 | 2.13E-08 | 0.0690 | 45.607 | %ID/g | 0.9845 | 3 |
| [22] | Breast | Intra-cranial | MCAIV | Albumin-Rho BSA | 7 | Passive | None | 6.84* | 1.80* | 1.90E-07* | 5.00E-08* | N/A | N/A | N/A | N/A | N/A |
| [22] | Breast | Intra-cranial | MCAIV | Albumin-Rho BSA | 7 | Passive | None | 10.44* | 5.40* | 2.90E-07* | 1.50E-07* | N/A | N/A | N/A | N/A | N/A |
| [11] | Pan-creatic | SubQ | BxPC3 | PEG-b-P(Glu) BCP Micelles | 30 | Passive | None | 1.334 | 0.176 | 3.71E-08 | 4.89E-09 | 0.0725 | 100 | %ID/g | 0.9582 | 4 |
| [11] | Pan-creatic | SubQ | BxPC3 | PEG-b-P(Glu) BCP Micelles | 50 | Passive | None | 0.872 | 0.107 | 2.42E-08 | 2.97E-09 | 0.0869 | 100 | %ID/g | 0.9276 | 4 |
| [11] | Pan-creatic | SubQ | BxPC3 | PEG-b-P(Glu) BCP Micelles | 70 | Passive | None | 0.541 | 0.096 | 1.50E-08 | 2.67E-09 | 0.0915 | 100 | %ID/g | 0.9147 | 4 |
| [11] | Pan-creatic | SubQ | BxPC3 | PEG-b-P(Glu) BCP Micelles | 90 | Passive | None | 0.579 | 0.081 | 1.61E-08 | 2.25E-09 | 0.1057 | 100 | %ID/g | 0.9029 | 4 |
| [7] | Colon | SubQ | C26 | PEG-CL liposome (targeted) | 100 | Active | Trans-ferrin | 1.187 | 0.354 | 3.30E-08 | 9.83E-09 | 0.0500 | 22.424 | %ID/g | 0.768 | 5 |
| [7] | Colon | SubQ | C26 | PEG-CL liposome (without targeting) | 100 | Passive | None | 0.982 | 0.391 | 2.73E-08 | 1.09E-08 | 0.0600 | 19.865 | %ID/g | 0.849 | 4 |
| [23] | Colon | SubQ | C26 | Liposome vinorelbine (VNBL) | 102 ± 6.9 | Passive | None | 5.611 | 4.185 | 1.56E-07 | 1.16E-07 | 0.0350 | 37.623 | %IA/g | 0.8133 | 5 |
| [23] | Colon | SubQ | C26 | Liposome Dox (DXRL) | 131 ± 30 | Passive | None | 1.552 | 0.581 | 4.31E-08 | 1.61E-08 | 0.0770 | 27.093 | %IA/g | 0.9669 | 5 |
| [24] | Colon | SubQ | LS174T | Albumin-Rho BSA | 7 | Passive | None | 4.32* | 1.80* | 1.20E-07* | 5.00E-08* | N/A | N/A | N/A | N/A | N/A |
| [24] | Colon | SubQ | LS174T | PEG Liposome | 100 | Passive | None | 0.72* | 0.58* | 2.00E-08* | 1.60E-08* | N/A | N/A | N/A | N/A | N/A |
| OD | Colon | SubQ | HT29 | 3HM | 20 | Passive | None | 1.531 | 0.327 | 4.25E-08 | 9.08E-09 | 0.0420 | 0.029 | mg/mL | 0.9986 | 4 |
| OD | Colon | SubQ | HT29 | Micelle | 20 | Passive | None | 2.704 | 1.14 | 7.51E-08 | 3.17E-08 | 0.2250 | 0.035 | mg/mL | 0.9998 | 4 |
| OD | Colon | SubQ | HT29 | PEG Liposome | 100 | Passive | None | 2.050 | 0.486 | 5.69E-08 | 1.35E-08 | 0.0690 | 0.0309 | mg/mL | 0.9618 | 4 |
| [11] | Colon | SubQ | C26 | PEG-b-P(Glu) BCP Micelles | 30 | Passive | None | 1.087 | 0.067 | 3.02E-08 | 1.86E-09 | 0.0725 | 100 | %ID/g | 0.9582 | 4 |
| [11] | Colon | SubQ | C26 | PEG-b-P(Glu) BCP Micelles | 50 | Passive | None | 1.234 | 0.165 | 3.43E-08 | 4.58E-09 | 0.0869 | 100 | %ID/g | 0.9276 | 4 |
| [11] | Colon | SubQ | C26 | PEG-b-P(Glu) BCP Micelles | 70 | Passive | None | 1.420 | 0.184 | 3.94E-08 | 5.11E-09 | 0.0915 | 100 | %ID/g | 0.9147 | 4 |
| [11] | Colon | SubQ | C26 | PEG-b-P(Glu) BCP Micelles | 90 | Passive | None | 1.139 | 0.075 | 3.16E-08 | 2.08E-09 | 0.1057 | 100 | %ID/g | 0.9029 | 4 |
| [22] | GBM | Ortho-topic, Intra-cranial | U87MG | Albumin-Rho BSA | 7 | Passive | None | 13.68* | 4.32* | 3.80E-07* | 1.20E-07* | N/A | N/A | N/A | N/A | N/A |
| [14] | GBM | Ortho-topic, Intra-cranial | U87MG | 3HM | 20 | Passive | None | 0.832 | 0.065 | 2.31E-08 | 1.81E-09 | 0.0446 | 0.272 | mg/mL | 0.8167 | 4 |
| [14] | GBM | Ortho-topic, Intra-cranial | U87MG | PEG Liposome | 100 | Passive | None | 0.389 | 0.010 | 1.08E-08 | 2.78E-10 | 0.0421 | 0.258 | mg/mL | 0.833 | 4 |
| [25] | GBM | Ortho-topic, Intra-cranial | 9L | PAMAM dendrimer | ~4 | Passive | None | 0.157** | 0.066 | 4.36E-09** | 1.83E-09 | 0.0880 | 30.073 | ug/g | 0.9452 | 6 |
| [26] | GBM | Ortho-topic, Intra-cranial | U87MG | NK012 polymeric micelle | ~20 | Passive | None | 1.156 | 0.389 | 3.21E-08 | 1.08E-08 | 0.0735 | 1096 | ng/g | 0.9586 | 4 |
| [27] | GBM | SubQ | U87MG | Single walled carbon nanotubes (PEG5400) | 1-5 nm length, 100-300 nm diam. | Passive | None | 7.301** | 2.269 | 2.03E-07** | 6.30E-08 | 0.1610 | 19.335 | %ID/g | 0.8019 | 5 |
| [28] | GBM | SubQ | U87MG | Nano Gold Tripods | 20.6 ± 0.5 | Passive | None | 2.257 | 1.416 | 6.27E-08 | 3.93E-08 | 0.0270 | 9.393 | %ID/g | 0.8931 | 4 |
| [28] | GBM | SubQ | U87MG | Nano Gold Tripods-RGD | 22.8 ± 0.6 | Active | RGD | 16.34** | 7.70 | 4.54E-07** | 2.14E-07 | 0.0440 | 17.901 | %ID/g | 0.9799 | 4 |
| [28] | GBM | SubQ | U87MG | Nano Gold Tripods-RGD Blocked (Control) | 22.8 ± 0.6 | Active | RGD (Blocked) | 2.656 | 1.358 | 7.38E-08 | 3.77E-08 | 0.0410 | 17.117 | %ID/g | 0.9146 | 4 |
| [29] | Skin | Dorsal skin fold window chamber | FaDu | Dextran | 3.3 kDa (3nm) | Passive | None | 9102 | 2296 | 2.53E-04 | 6.38E-05 | 4.3850 | 73.174 | %Vmax | 0.9445 | 12 |
| [29] | Skin | Dorsal skin fold window chamber | FaDu | Dextran | 10 kDa | Passive | None | 526.5 | 136.3 | 1.46E-05 | 3.79E-06 | 4.3940 | 82.483 | %Vmax | 0.9734 | 12 |
| [29] | Skin | Dorsal skin fold window chamber | FaDu | Dextran | 40 kDa | Passive | None | 236.73 | 18.24 | 6.58E-06 | 5.07E-07 | 2.1980 | 84.084 | %Vmax | 0.9205 | 12 |
| [29] | Skin | Dorsal skin fold window chamber | FaDu | Dextran | 70 kDa (14nm) | Passive | None | 202.0 | 21.32 | 5.61E-06 | 5.92E-07 | 1.8870 | 94.506 | %Vmax | 0.9865 | 12 |
| [29] | Skin | Dorsal skin fold window chamber | FaDu | Dextran | 2 MDa (15 nm) | Passive | None | 34.91 | 0.759 | 9.70E-07 | 2.11E-08 | 1.1650 | 99.14 | %Vmax | 0.9917 | 12 |
| [30] | NSCLC | SubQ | H520 | PEG Liposome (PEG2000) | 81.9 ±3.7 | Passive | None | 0.861 | 0.138 | 2.39E-08 | 3.83E-09 | 0.0259 | 5.319 | mg/mL | 0.9535 | 7 |
| [30] | NSCLC | SubQ | H520 | PEG Liposome-0.64%mol NGR (PEG2000) | 84.1 ±3.5 | Active | NGR | 2.201 | 0.214 | 6.11E-08 | 5.94E-09 | 0.0284 | 5.817 | mg/mL | 0.9692 | 7 |
| [30] | NSCLC | SubQ | H520 | PEG Liposome-2.56%mol NGR (PEG2000) | 86.8±2.7 | Active | NGR | 1.468 | 0.119 | 4.08E-08 | 3.31E-09 | 0.0308 | 6.329 | mg/mL | 0.9661 | 7 |
| [30] | NSCLC | SubQ | H520 | PEG Liposome-0.64%mol NGR (PEG3400) | 86.7±2.0 | Active | NGR | 1.800** | 0.651 | 5.00E-08** | 1.81E-08 | 0.0291 | 5.203 | mg/mL | 0.9608 | 7 |
| [30] | NSCLC | SubQ | H520 | PEG Liposome-0.64%mol ARA (PEG2000) (control) | Not reported | Passive | ARA (mis-match) | 1.111 | 0.207 | 3.09E-08 | 5.75E-09 | 0.0278 | 4.4828 | mg/mL | 0.8843 | 7 |
| [30] | NSCLC | SubQ | H520 | PEG Liposome-0.64%mol ARA (PEG3400) (control) | Not reported | Passive | ARA (mis-match) | 0.971 | 0.062 | 2.70E-08 | 1.72E-09 | 0.0234 | 4.3885 | mg/mL | 0.9537 | 7 |

### References

[1]	H. Hong, Y. Zhang, J.W. Engle, T.R. Nayak, C.P. Theuer, R.J. Nickles, T.E. Barnhart, W. Cai, Biomaterials In vivo targeting and positron emission tomography imaging of tumor vasculature with 66 Ga-labeled nano-graphene, Biomaterials. 33 (2012) 4147–4156. doi:10.1016/j.biomaterials.2012.02.031.

[2]	J. Guo, H. Hong, G. Chen, S. Shi, Q. Zheng, Y. Zhang, C.P. Theuer, T.E. Barnhart, W. Cai, S. Gong, Image-guided and tumor-targeted drug delivery with radiolabeled unimolecular micelles, Biomaterials. 34 (2013) 8323–8332. doi:10.1016/j.biomaterials.2013.07.085.

[3]	H. Lee, B. Hoang, H. Fonge, R.M. Reilly, C. Allen, In vivo distribution of polymeric nanoparticles at the whole-body, tumor, and cellular levels, Pharm. Res. 27 (2010) 2343–2355. doi:10.1007/s11095-010-0068-z.

[4]	H. Lee, H. Fonge, B. Hoang, R.M. Reilly, C. Allen, The effects of particle size and molecular targeting on the intratumoral and subcellular distribution of polymeric nanoparticles, Mol. Pharm. 7 (2010) 1195–1208. doi:10.1021/mp100038h.

[5]	X. Guo, C. Shi, J. Wang, S. Di, S. Zhou, PH-triggered intracellular release from actively targeting polymer micelles, Biomaterials. 34 (2013) 4544–4554. doi:10.1016/j.biomaterials.2013.02.071.

[6]	Y. Ma, M. Sadoqi, J. Shao, Biodistribution of indocyanine green-loaded nanoparticles with surface modifications of PEG and folic acid, Int. J. Pharm. 436 (2012) 25–31. doi:10.1016/j.ijpharm.2012.06.007.

[7]	Y. Miyajima, H. Nakamura, Y. Kuwata, J. Lee, S. Masunaga, K. Ono, K. Maruyama, Transferrin-Loaded nido -Carborane Liposomes : Tumor-Targeting Boron Delivery System for Neutron Capture Therapy, Bioconjug. Chem. 17 (2006) 1314–1320. doi:10.1021/bc060064k.

[8]	M. Dunne, J. Zheng, J. Rosenblat, D.A. Jaffray, C. Allen, APN / CD13-targeting as a strategy to alter the tumor accumulation of liposomes, J. Control. Release. 154 (2011) 298–305. doi:10.1016/j.jconrel.2011.05.022.

[9]	T. Lammers, F. Kiessling, W.E. Hennink, G. Storm, Drug targeting to tumors: Principles, pitfalls and (pre-) clinical progress, J. Control. Release. 161 (2012) 175–187. doi:10.1016/j.jconrel.2011.09.063.

[10]	M. Juweid, R. Neumann, C. Paik, M.J. Perez-Bacete, J. Sato, W. van Osdol, J.N. Weinstein, Micropharmacology of Monoclonal Antibodies in Solid Tumors: Direct Experimental Evidence for a Binding Site Barrier, Cancer Res. 52 (1992) 5144–5153.

[11]	H. Cabral, Y. Matsumoto, K. Mizuno, Q. Chen, M. Murakami, M. Kimura, Y. Terada, M.R. Kano, K. Miyazono, M. Uesaka, N. Nishiyama, K. Kataoka, Accumulation of sub-100 nm polymeric micelles in poorly permeable tumours depends on size, Nat. Nanotechnol. 6 (2011) 815–823. doi:10.1038/nnano.2011.166.

[12]	C. Wong, T. Stylianopoulos, J. Cui, J. Martin, V.P. Chauhan, W. Jiang, Z. Popovic, R.K. Jain, M.G. Bawendi, D. Fukumura, Multistage nanoparticle delivery system for deep penetration into tumor tissue, PNAS. 108 (2011) 2426–2431. doi:10.1073/pnas.1018382108.

[13]	H.-J. Li, J.-Z. Du, X.-J. Du, C.-F. Xu, C.-Y. Sun, H.-X. Wang, Z.-T. Cao, X.-Z. Yang, Y.-H. Zhu, S. Nie, J. Wang, Stimuli-responsive clustered nanoparticles for improved tumor penetration and therapeutic efficacy, Proc. Natl. Acad. Sci. 113 (2016) 4164–4169. doi:10.1073/pnas.1522080113.

[14]	J.W. Seo, J. Ang, L.M. Mahakian, S. Tam, B. Fite, E.S. Ingham, J. Beyer, J. Forsayeth, K.S. Bankiewicz, T. Xu, K.W. Ferrara, Self-assembled 20-nm 64Cu-micelles enhance accumulation in rat glioblastoma, J. Control. Release. 220 (2015) 51–60. doi:10.1016/j.jconrel.2015.09.057.

[15]	R.T. Tong, Y. Boucher, S. V Kozin, F. Winkler, D.J. Hicklin, R.K. Jain, Vascular Normalization by Vascular Endothelial Growth Factor Receptor 2 Blockade Induces a Pressure Gradient Across the Vasculature and Improves Drug Penetration in Tumors, Cancer Res. 64 (2004) 3731–3736.

[16]	N.Z. Wu, D. Da, T.L. Rudoll, S. Liposomes, D. Needham, A.R. Whorton, M.W. Dewhirst, Increased Microvascular Permeability Contributes to Preferential Accumulation of Stealth Liposomes in Tumor Tissue Increased Microvascular Permeability Contributes to Preferential Accumulation of, Cancer Res. 53 (1993) 3765–3770.

[17]	K.C.L. Black, Y. Wang, H.P. Luehmann, X. Cai, W. Xing, B. Pang, Y. Zhao, C.S. Cutler, L. V. Wang, Y. Liu, Y. Xia, Radioactive198Au-doped nanostructures with different shapes for in vivo analyses of their biodistribution, tumor uptake, and intratumoral distribution, ACS Nano. 8 (2014) 4385–4394. doi:10.1021/nn406258m.

[18]	A. Pathak, P. Kumar, K. Chuttani, S. Jain, A.K. Mishra, S.P. Vyas, K.C. Gupta, Gene Expression, Biodistribution, and Pharmacoscintigraphic Evaluation of Chondroitin Sulfate-PEI Nanoconstructs Mediated Tumor Gene Therapy, ACS Nano. 3 (2009) 1493–1505.

[19]	A.W. Wong, E. Ormsby, H. Zhang, J.W. Seo, L.M. Mahakian, C.F. Caskey, K.W. Ferrara, A comparison of image contrast with 64Cu-labeled long circulating liposomes and 18F-FDG in a murine model of mammary carcinoma, Am. J. Nucl. Med. Mol. Imaging. 3 (2013) 32–43. doi:10.2522/ptj.20100142.

[20]	N. Dube, J.W. Seo, H. Dong, J.Y. Shu, R. Lund, L.M. Mahakian, K.W. Ferrara, T. Xu, Effect of Alkyl Length of Peptide − Polymer Amphiphile on Cargo Encapsulation Stability and Pharmacokinetics of 3 ‑ Helix Micelles, Biomacromolecules. 15 (2014) 2963–2970.

[21]	C. Pérez-Medina, D. Abdel-Atti, Y. Zhang, V.A. Longo, C.P. Irwin, T. Binderup, J. Ruiz-Cabello, Z.A. Fayad, J.S. Lewis, W.J.M. Mulder, T. Reiner, A Modular Labeling Strategy for In Vivo PET and Near-Infrared Fluorescence Imaging of Nanoparticle Tumor Targeting, J. Nucl. Med. 55 (2014) 1706–1711. doi:10.2967/jnumed.114.141861.

[22]	F. Yuan, H.A. Salehi, Y. Boucher, R.K. Jain, U.S. Vasthare, R.F. Tuma, Vascular Permeability and Microcirculation of Gliomas and Mammary Carcinomas Transplanted in Rat and Mouse Cranial Windows, Cancer Res. 54 (1994) 4564–4568.

[23]	Y.-Y. Lin, C.-H. Chang, J.-J. Li, M.G. Stabin, Y.-J. Chang, L.-C. Chen, M.-H. Lin, Y.-L. Tseng, W.-J. Lin, T.-W. Lee, G. Ting, C.A. Chang, F.-D. Chen, H.-E. Wang, Pharmacokinetics and Dosimetry of ^111^ In/ ^188^ Re-Labeled PEGylated Liposomal Drugs in Two Colon Carcinoma-Bearing Mouse Models, Cancer Biother. Radiopharm. 26 (2011) 373–380. doi:10.1089/cbr.2010.0906.

[24]	F. Yuan, M. Leunig, S.K. Huang, D.A. Berk, D. Papahadjopoulos, R.K. Jam, Microvascular Permeability and Interstitial Penetration of Sterically Stabilized ( Stealth ) Liposomes in a Human Tumor Xenograft ’, Cancer Res. 54 (1994) 3352–3357.

[25]	F. Zhang, P. Mastorakos, M.K. Mishra, A. Mangraviti, J. Zhou, J. Hanes, H. Brem, A. Olivi, R. Kannan, Uniform brain tumor distribution and tumor associated macrophage targeting of systemically administered dendrimers, Biomaterials. 52 (2015) 507–516. doi:10.1016/j.biomaterials.2015.02.053.Uniform.

[26]	J.I. Kuroda, J.I. Kuratsu, M. Yasunaga, Y. Koga, Y. Saito, Y. Matsumura, Potent antitumor effect of SN-38-incorporating polymeric micelle, NK012, against malignant glioma, Int. J. Cancer. 124 (2009) 2505–2511. doi:10.1002/ijc.24171.

[27]	Z. Liu, W. Cai, L. He, N. Nakayama, K. Chen, X. Sun, X. Chen, H. Dai, In vivo biodistribution and highly efficient tumour targeting of carbon nanotubes in mice, Nat. Nanotechnol. 2 (2007) 47–52. doi:10.1038/nnano.2006.170.

[28]	K. Cheng, S.R. Kothapalli, H. Liu, A.L. Koh, J. V. Jokerst, H. Jiang, M. Yang, J. Li, J. Levi, J.C. Wu, S.S. Gambhir, Z. Cheng, Construction and validation of nano gold tripods for molecular imaging of living subjects, J. Am. Chem. Soc. 136 (2014) 3560–3571. doi:10.1021/ja412001e.

[29]	M.R. Dreher, W. Liu, C.R. Michelich, M.W. Dewhirst, F. Yuan, A. Chilkoti, Tumor vascular permeability, accumulation, and penetration of macromolecular drug carriers, J. Natl. Cancer Inst. 98 (2006) 335–344. doi:10.1093/jnci/djj070.

[30]	J. Zheng, J. Liu, M. Dunne, D.A. Jaffray, C. Allen, In Vivo Performance of a Liposomal Vascular Contrast Agent for CT and MR-Based Image Guidance Applications, 24 (2007). doi:10.1007/s11095-006-9220-1.
